# Duck, duck, goose: Benchmark bird surveys help quantify counting errors and bias in a citizen-science database

**DOI:** 10.1101/2020.12.09.418145

**Authors:** W. Douglas Robinson, Tyler A. Hallman, Rebecca A. Hutchinson

**Author notes:** **Correspondence:** W. Douglas Robinson.

## Abstract

The growth of biodiversity data sets generated by citizen scientists continues to accelerate. The availability of such data has greatly expanded the scale of questions researchers can address. Yet, error, bias, and noise continue to be serious concerns for analysts, particularly when data being contributed to these giant online data sets are difficult to verify. Counts of birds contributed to eBird, the world’s largest biodiversity online database, present a potentially useful resource for tracking trends over time and space in species’ abundances. We quantified counting errors in a sample of 1406 eBird checklists by comparing numbers contributed by birders (N=246) who visited a popular birding location in Oregon, USA, with numbers generated by a professional ornithologist engaged in a long-term study creating benchmark (reference) measurements of daily waterbird counts. We focused on waterbirds, which are easily visible at this site. We evaluated potential predictors of count differences, including characteristics of contributed checklists, of each species, and of time of day and year. Count differences were biased toward undercounts, with more than 75% of counts being below the daily benchmark value. When only checklists that actually reported a species known to be present were included, median count errors were −29.1% (range: 0 to −42.8 %; N=20 species). Model sets revealed an important influence of each species’ reference count, which varied seasonally as waterbird numbers fluctuated, and of percent of species known to be present each day that were included on each checklist. That is, checklists indicating a more thorough survey of the species richness at the site also had, on average, lower counting errors. However, even on checklists with the most thorough species lists, counts were biased low and exceptionally variable in their accuracy. To improve utility of such bird count data, we suggest three strategies to pursue in the future. One is to assess additional options for analytically determining how to select checklists that have the highest probability of including less biased count data, as well as exploring options for correcting bias during the analysis stage. Another is to add options for users to provide additional information that helps analysts choose checklists, such as an option for users to tag checklists where they focused on obtaining accurate counts. We also recommend exploration of opportunities to effectively calibrate citizen-science bird count data by establishing a formalized network of marquis sites where dedicated observers regularly contribute carefully collected benchmark data.

## Introduction

Contributions of volunteers to scientific databases are increasing as the popularity of citizen science continues to grow (Miller-Rushing et al., 2012; Chandler et al., 2017). Many citizen science projects are open-access and anyone can contribute observations without required training in best data collection practices (Cohn, 2008). eBird is an open online database with more than 560,000 users (eBirders) contributing millions of bird observations annually via checklists (Sullivan et al., 2009). Each checklist contains a list of bird species identified on a particular date and, ideally, counts of each species, as well as information on location visited, basic protocol used while birding (traveling, staying stationary, etc.), and duration of effort (Wood et al., 2011). The huge spatial extent of presence-absence data in eBird has facilitated efforts to model species distributions across continental and global spatial scales once data have been filtered to exclude potentially problematic checklists (Fink et al., 2013). The degree to which the count data may reliably inform scientific and management objectives remains unclear.

Although efforts to quantify issues associated with bird species detection have been studied and continue to be developed, both in citizen science databases and in structured scientific surveys (Buckland et al., 2008; Hutto, 2016; Walker and Taylor, 2017), less is known about potential counting errors and biases leading to noisy data. Counting birds is difficult, even by the most proficient observers (Robbins and Stallcup, 1981; Robinson et al., 2018). Methods to account for detection issues in bird counting studies continue to expand with development of new data collection and analytical methods (Buckland et al., 2008; Barker et al., 2018). Nearly all the methods, however, require a sophisticated sampling protocol that would exclude most volunteer birder contributions and therefore limit the advantages of gathering data at massive geographic scales. Yet, the potential windfall from large quantities of data can quickly be eroded if a lack of structured protocols leads to data quality concerns (Kelling et al., 2019). Given that abundance is one of the fundamental influences on population dynamics, functional roles in ecosystems, and even extinction risk (Brown, 1984), a better understanding of the potential value of count data contributed to massive online databases by untrained volunteers is needed (Greenwood, 2007). For example, species count errors in eBird data could limit our abilities to observe important abundance trends (Horns et al., 2018). Effective processes for evaluating and handling such errors need further development, owing to the potentially huge value of tracking population changes at a continental scale during this era of rapid environmental change (Bird et al., 2014; Fink et al., 2020).

Among the primary concerns are errors, bias and noise. Errors, for our purposes here, are differences in counts between a reference (benchmark) value and values included in eBird checklists for the same species on the same date. Errors are comprised of both bias and noise. Bias is the tendency for the errors to be consistently higher or lower than the reference value. Noise is the additional random counting error that increases variance of the counts. All three impede efforts to determine true count values, and are challenges common to many branches of biology (West, 1999; Guillery, 2002). We acknowledge that labeling such count differences as errors assumes the benchmark values have less error and doing so risks offending some eBird contributors. Given that reliable benchmarks are achieved by consistent application of best counting practices, we do consider deviations from the benchmark values here to be errors, not simply variation among observers. To acknowledge that there are sources of error in all measurements, however, we often refer to such deviations as count differences. We consider the terms ‘error’ and ‘count differences’ to be synonymous.

Robust comparisons of count differences are improved when data are collected in situations where detectability challenges are expected to be low. Such situations are rare but uniquely valuable. We used an extensive data set focused on benchmarking the richness and abundances of birds at a water treatment site in Oregon, USA. We compared count data gathered by a professional ornithologist focused specifically on creating an accurate benchmark measurement of daily fluctuations in waterbird counts with counts submitted by birders to eBird. We quantified the magnitude and directionality of count differences. Our data span 10 years and include 1406 eBird checklists contributed by 246 observers, as well as 2038 checklists in the benchmark data. The site is well suited for rigorous comparisons because all waterbirds are in the open, largely tolerant of human activity, and so provide a best-case scenario for detection, identification, and counting of birds. No adjustments for detectability or availability issues should be needed because all parts of the ponds are visible. Thus, discrepancies in counts between a professional observer focused on obtaining accurate numbers and data reported to eBird should be attributable to counting errors instead of availability and detectability issues. While there could be very minor detectability issues, like some diving waterbirds being under water briefly, the vast majority of error in this setting should be attributable to counting error.

We first quantified count differences then sought to understand potential factors explaining the magnitude and directionality of count differences. We hypothesized that counting errors would be influenced by traits associated with the species being counted, with an index of observer experience (percent of species detected), and with seasonal changes in numbers of birds present. For example, we expected count differences might be slightly greater for diving ducks, which are sometimes briefly under water while foraging, and lower for dabbling species, which sit in the open continuously. We expected smaller count differences in checklists that included a higher proportion of the species present each day. We also hypothesized that count differences would be greater when overall total number of waterbirds present was high, potentially causing observers to be overwhelmed and therefore more prone to counting errors. Finally, we explored the possibility that, even if count data were biased on individual checklists, the waterbird community might be adequately characterized as a whole by combining count data from multiple observers and checklists. We conclude by proposing additional approaches that may reveal the extent to which citizen-science bird count data may be used to estimate abundances reliably.

## Methods

### Study Area

Bird count data were gathered from 2010 to 2019 at the Philomath Wastewater Treatment facility, in Philomath, Oregon USA. The site contained two 35-ha ponds until 2011 when two additional 35-ha ponds were added. Each pond is rectangular and enclosed by a berm with a single-lane road. Birders circumnavigate the ponds typically by vehicle, rarely by walking or bicycling; WDR drove. Vegetation does not obscure the view at any pond. All shores are covered by large rocks (riprap). Birders circle all four ponds during a visit, very rarely restricting visits to fewer ponds. We found that the distribution of visit durations was unimodal (median = 60 min; Median Average Deviation (MAD) = 37; skew=1.161; N=1646 checklists) suggesting that birders use similar methods while at the ponds.

### Study species

We included 20 species we refer to as “waterbirds,” species that swim in the open while on the ponds and should be easily seen (Table 1). The species are primarily ducks and geese, but also include grebes, American Coot (*Fulica americana*), and gulls. These are species birders identify by sight, not by sound. We excluded species that occurred primarily as fly-overs, such as Cackling Goose (*Branta hutchinsi*), species whose counts rarely exceeded two per day, and species whose numbers varied strongly within a day. The number of waterbirds present at the site varied seasonally from a few dozen during mid-summer (June) to five thousand or more during fall migration (October-November).

**Table 1.** Twenty species were included in the study. Scientific names, sequence, and short-hand codes follow American Ornithological Society (http://checklist.aou.org/taxa). See text for definitions of dabbler versus diver and dispersed versus aggregated foragers. Plumage sexual dichromatism was scored based on the period of year in which the species is most numerous at the study site: weak or no dichromatism (0) and moderate to strong dichromatism (1).

| English name | Scientific name | Code | Dabbler (0) or Diver (1) | Dispersed (0) or aggregated (1) | Plumage dichromatism |
| --- | --- | --- | --- | --- | --- |
| **Wood Duck** | *Aix sponsa* | wodu | 0 | 0 | 1 |
| **Cinnamon Teal** | *Spatula cyanoptera* | cite | 0 | 0 | 0 |
| **Northern Shoveler** | *Spatula clypeata* | nsho | 0 | 1 | 1 |
| **Gadwall** | *Mareca strepera* | gadw | 0 | 0 | 1 |
| **American Wigeon** | *Mareca americana* | amwi | 0 | 1 | 1 |
| **Mallard** | *Anas platyrhynchos* | mall | 0 | 0 | 1 |
| **Northern Pintail** | *Anas acuta* | nopi | 0 | 0 | 1 |
| **Green-winged Teal** | *Anas crecca* | gwte | 0 | 1 | 1 |
| **Canvasback** | *Aythya valisineria* | canv | 1 | 0 | 1 |
| **Ring-necked Duck** | *Aythya collaris* | rndu | 1 | 1 | 1 |
| **Lesser Scaup** | *Aythya affinis* | lesc | 1 | 0 | 1 |
| **Surf Scoter** | *Melanitta perspicillata* | susc | 1 | 0 | 0 |
| **Bufflehead** | *Bucephala albeola* | buff | 1 | 0 | 1 |
| **Hooded Merganser** | *Lophodytes cucullatus* | home | 1 | 0 | 0 |
| **Ruddy Duck** | *Oxyura jamaicensis* | rudu | 1 | 1 | 0 |
| **Pied-billed Grebe** | *Podilymbus podiceps* | pbgr | 1 | 0 | 0 |
| **Eared Grebe** | *Podiceps nigricollis* | eagr | 1 | 0 | 0 |
| **American Coot** | *Fulica americana* | amco | 0 | 1 | 0 |
| **Ring-billed Gull** | *Larus delawarensis* | rbgu | 0 | 0 | 0 |
| **California Gull** | *Larus californicus* | cagu | 0 | 0 | 0 |

### Benchmark counts

All birds of all species were counted during each site visit by WDR. We call these our benchmark counts (*R*\*) and they serve as the reference values against which all other count data are compared. Waterbird counts were made to plus or minus one individual except for Northern Shoveler (*Spatula clypeata*), which were plus or minus 10 because they forage in constantly moving dense aggregations rendering more precise counts problematic, and Bufflehead (*Bucephala albeola*), which were counted to plus or minus 5 because they dive so frequently while foraging in the early morning period surveyed by WDR that more accurate counts were difficult. Counts were tallied separately for each pond then aggregated later. In the time frame of these counts, movements between ponds were normally minimal. Duration of counting time was recorded separately for each pond. To reduce possible use of WDR’s count data by eBirders who wanted to post numbers but may not have counted on their own, we used three steps to minimize copying of data. First, we imposed a time lag of one to four weeks between dates of counting and of uploading to eBird the WDR data. Second, we hid all of WDR’s checklists from the eBird public display in Recent Visits and, third, we posted only the pond-specific data, not the aggregated data. We used aggregated counts from the first visit each day as *R*\* for comparison with counts reported on eBird checklists.

On some days (N=84), WDR counted birds more than once. These second-visit data, which we call Ref2 counts, were also complete counts of the study species and averaged 13% shorter in duration, yet counts were generally similar. They were used to characterize within-day variability in numbers but provide a conservative estimate of that variability because they were largely conducted on days with exceptional levels of migratory movements. Thus, they estimate a probable upper bound on the expected amount of within-day variability in waterbird numbers (averaging 0 to −8%). We also used these Ref2 data to evaluate of time-of-day effects when comparing WDR counts with data from the ten observers contributing the most study site data to eBird, because eBirders tended to count birds later in the day than did WDR. The times of day eBird checklists were initiated as well as the difference in start times of eBird and benchmark checklists were unimportant in predicting percent error in our across-species and species-specific model sets. Therefore, we concluded that comparisons of count differences between *R*\* and eBird checklists were appropriate and that possible time-of-day effects could be ignored.

### eBird checklists

We downloaded eBird checklists from the Philomath Sewage Ponds eBird hotspot as well as eBirder personal locations within 1 km from 2010 to 2019. Only data obviously restricted to the ponds were included. No other waterbird sites are present within 4 km of the site. Most eBirders used the pre-established hotspot as the checklist location but some created new personal locations each time. We included eBird checklists following the stationary, traveling, and area protocols. We removed checklists with greater than ten observers or durations of over five hours. We included only complete checklists with all birds reported and removed any checklists where observers reported no waterbirds. From each complete eBird checklist, we collected data on date, start time, observer, duration of count, identity of waterbird species reported (to allow calculation of percent richness; see below), and count data for our twenty focal species. When species were recorded as present but not counted (X noted instead of a number), those data were excluded because no count difference could be calculated.

### Comparisons of count data

We restricted our comparisons to dates where WDR counted birds and at least one eBird checklist was contributed on the same day (N=767 dates). Our questions were about counting differences and not detection rates of rare species, so we further restricted our comparisons to counts of greater than three for each species detected on WDR’s first visit (*R*\*). We calculated the *Count Difference* for each species by subtracting *R*\* from eBird counts on each checklist. Count differences were positive when eBird checklists reported higher numbers than *R*\* or negative when eBird checklists reported fewer birds than *R*\*. Numeric values of count differences spanned three orders of magnitude, so we focus on reporting *Percent Error*, which we calculated by converting each difference to a proportion of *R*\*.

### Hypothesized predictors of percent error

To evaluate factors hypothesized to be associated with percent error, we included variables associated with species, checklists, time of year and observer experience. *Species characteristics* included categorization as dabbler versus diver, degree to which species form dense aggregations, and the degree of sexual dimorphism. *Checklist characteristics* included start time, duration and number of observers. *Time-of-year characteristics* were associated with daily numbers of waterbirds (*R*\*, Ref2 and their sums for all 20 species) and waterbird species richness present at the study site (measured as the richness detected by the professional [proRichness] as well as the aggregate of species listed in eBird checklists and proRichness). Because *observer experience* at the site might also influence counting accuracy, we compared data from the 10 observers who contributed the most checklists with the *R*\* and Ref2 benchmark data. Additional details on each variable are explained below.

#### Species characteristics

To explore patterns of species-specific variability in count data, we created categorical variables for species traits that might impact counts (Table 1). We categorized birds as dabblers versus divers. Dabblers were any species that foraged primarily by swimming on the surface of the water, which included gulls, American Coot, and *Aix*, *Anas*, *Mareca,* and *Spatula* ducks. Divers foraged below water regularly and included scoters, grebes, and *Aythya* and *Bucephala* ducks.

We also included an index of spatial aggregation on the ponds. Some species, for example Northern Shoveler, often forage in densely packed groups, creating challenging circumstances to accurately count birds, while other species forage singly or as spatially-distanced groups where enumeration should be much easier. The aggregation index was simply a subjective binary classification (0 for foraging alone or in loose aggregations versus 1 for foraging in aggregations that might render counting difficult) based on our years of experience at the site.

The degree of plumage dimorphism and similarity to other species could influence error and bias in counts because of species misidentification. We categorized species as those with weak or no obvious plumage dichromatism during most of the period of time when each species was present (e.g., geese, coots) versus strong dichromatism (males and females distinctly visually different).

To evaluate the possibility that species identification of similar species might influence count differences, we used another subjective binary category called “Doppelganger;” 1 indicated the species co-occurred with a similar species whereas 0 indicated the species was unique in appearance and unlikely to be confused with other species. The categorization may vary seasonally, especially in late summer when many waterbirds molt to eclipse plumage. Because total waterbird numbers were low during late summer, we utilized one value for each species.

#### Checklist characteristics

Daily start time among eBird checklists was highly variable, covering all daylight hours. The mean start time was 4 hours later than the mean start time for WDR visits. Although we only compared counts conducted on the same day, we wanted to evaluate potential effects of time-of-day and temporal lag between the eBird checklist counts and *R*\*. To do so, we converted checklist start time to minutes since midnight then calculated the difference in start time between eBird checklists and WDR first visits.

Because our Ref2 counts occurred later in the day when more eBird checklists were initiated, we included Ref2 as an “additional observer” in some comparisons to provide an important check on within-day variability in counts as a possible explanation for count differences between *R*\* and eBird checklists. Because Ref2 counts were generated on days with high levels of migratory movement, we consider the count differences between *R*\* and Ref2 to represent an upper bound on expected levels of within-day variability in waterbird numbers.

Additional factors associated with each checklist could influence count differences. We reasoned that duration of time spent at the site should be positively related to count accuracy. All complete eBird checklists are required to have a measurement of event duration.

Number of observers might also influence counting accuracy, so we included the reported number of observers for each eBird checklist. The *R*\* and Ref2 counts were made when WDR was alone more than 99% of all dates.

#### Time-of-year characteristics

Date influences the number of species present as well as the abundances of each species. Both richness and abundance could influence counting accuracy so we included day of year in our models. Because we hypothesized that total number of all waterbirds combined may influence counting accuracy, we included *R*\* counts of all 20 study species and the combined daily total of all waterbirds in our model sets. In that way, we established the baseline numbers of waterbirds known to be present as a function of date. In calculating total waterbird abundance, we used data limited to the 20 study species and excluded a subset of species known to have high daily variability in counts, such as geese, which occurred primarily as fly-overs. The other species excluded from our focal group of 20 species were numerically rare. Further, to determine if percent error was influenced by the number of each particular species as opposed to overall waterbird abundance, we included *R*\* of each relevant species in our model sets.

We hypothesized overall waterbird species richness present at the site on a given date may influence counting accuracy. A higher number of species to identify could reduce focus for achieving accurate counts, particularly for the more regularly-occurring and common species (e.g., Mallards, Northern Shovelers). Therefore, we included in our models the total waterbird richness detected by WDR each day. Our analyses indicated that richness observed by WDR and total waterbird richness detected by all eBird contributors were highly correlated. We calculated daily *Percent Richness* based on the 35 possible waterbird species at the site and included that richness in our models (see Supplemental Text for a list of species). The other 15 species that formed our set of 35 waterbird species included: Snow Goose (*Anser caerulescens*), Greater White-fronted Goose (*Anser albifrons*), Cackling Goose (*Branta hutchinsii*), Canada Goose (*Branta canadensis*), Blue-winged Teal (*Spatula discors*), Eurasian Wigeon (*Mareca penelope*), Redhead (*Aythya americana*), Tufted Duck (*Aythya fuligula*), Greater Scaup (*Aythya marila*), White-winged Scoter (*Melanitta deglandi*), Black Scoter (*Melanitta americana*), Long-tailed Duck (*Clangula hyemalis*), Common Goldeneye (*Bucephala clangula*), Barrow’s Goldeneye (*Bucephala islandica*), and Common Merganser (*Mergus merganser*).

#### Observer experience

Observer experience at the site could also be influential, so we compared percent error in counts from the ten observers contributing the most eBird checklists at our study site with the *R*\* and Ref2 counts.

### Data analyses

We used the “lmer” package in R (R Core Team, 2020) to run mixed-effects models. Our overarching goal was to identify factors informative for explaining variation in *Percent Error*, our dependent variable in all models. We included observer ID and species as random effects to account for observer- and species-specific error when appropriate. We included four categorical species characteristics as fixed effects in our model sets: Dabbler or Diver; Sexually Dichromatic or not; Doppelganger or not; and Aggregated or not. Five checklist-related characteristics were included as fixed effects: start time (minutes since midnight), difference in start time between WDR’s first count of a day and each eBird checklist, duration (minutes), number of observers, and day of year. Four fixed-effects related to time-of-year were also included: *R*\* (WDR’s reference count of each species, which varied seasonally), waterbird abundance (aggregated across all species), total waterbird species richness and percent richness, our index of observer skill at species identification. We included models with the quadratic effects of species-specific abundance, waterbird abundance, waterbird richness, duration, number of observers, day of year, and percent richness to examine potential nonlinear shapes of their effects.

Before running mixed effects models, we scaled and centered all numeric variables. We assessed model performance through BIC and propagated best-performing shapes for each variable to multi-variable models. We used a forward stepwise approach and added additional potentially influential variables to the best-performing model until a stable (i.e., model remained the top model after the inclusion of additional variables) top-performing BIC model was identified.

Although *count difference* was normally distributed, *percent error* was not. Non-detections of species that were detected by WDR (eBird counts of zero) equal negative 100 *percent error*. Non-detections caused a bimodal distribution of *percent error* with a second peak at negative 100 percent. We removed non-detections to create a unimodal distribution of percent error. When non-detections were removed, *percent error* was heavily right-skewed due to the high number of negative *percent errors* and the few very large positive *percent errors*. To adjust skew, we added a constant to make all values positive and log (base 10) transformed percent error. In addition to adjusting skew, removal of non-detections improved the focus of our analyses on count errors, reducing chances that inclusion of zero counts of species might actually be species detection or identification problems instead of counting errors. Our restriction of counting error analyses to species detected in numbers of 3 or greater probably limited most effects of zero counts. In this paper we focus on analyses of data excluding non-detections but report some analyses in supplemental materials to show the effects of including non-detections (zero counts) on results. It is possible that an unknown number of zero counts were a result of reporting errors (data entry mistakes), but we assume this type of error is relatively rare.

#### Species-specific model sets

To understand the (in)consistency of variables influencing species-specific percent error, we ran standardized linear model sets of the effects of the explanatory variables described above on *transformed percent error* for each species. As above, we included models with quadratic effects of species abundance, waterbird abundance, waterbird richness, duration, number of observers, day of year, and percent richness. As each model set was species-specific, we excluded variables of species characteristics from these model sets. We included observer ID as an explanatory variable to examine its comparative influence. In these standardized model sets, we included separate models of the main effect of each variable and propagated the best shape for each variable into more complex models. Since start time and difference in start time were highly correlated, we use the top-performing of the two in subsequent models. We used a forward step-wise approach to determine the top-performing model of checklist covariates. We then ran models with pairs of all non-checklist explanatory variables with and without the variables in the top checklist covariate model. We used BIC to compare model performance and select top models.

#### Non-metric Multidimensional Scaling (NMDS)

To compare the overall communities described in eBird checklists, we conducted ordination in species space with NMDS on count data. We grouped checklists by observers to simplify the analysis. To visualize differences in community characterization, we chose to contrast January and October because January represents a time of year when waterbird migration is minimal and so daily numbers are relatively stable, whereas migration is at its peak during October, so richness is high and volatility in numbers can be high. To evaluate how characterization of waterbird abundance at these times varied with respect to eBirder checklists, we first removed all checklists that included an “X” for the count of any of our 20 study species. We then calculated the mean and median values of species counts across checklists for each observer during each month. To evaluate the idea that group collective contributions of multiple eBird checklists might characterize the waterbird community more similarly to *R*\*, we calculated mean counts of species across observers in January and October to create combined count values, which we call the Borg number 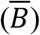. We similarly aggregated WDR’s first-visit species counts as a Reference community. To ensure that our 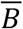 NMDS positions in species space were not driven overwhelmingly by an eBirder with the largest number of checklists, we reran the NMDS without checklists from the top-contributing observer included in 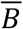. We used two dimensions and a maximum of 20 iterations to run NMDS with the “vegan” package in R (version 3.6.1).

## Results

We compared benchmark counts of waterbirds from WDR (*R*\*) and at least one eBirder on 672 dates, representing a total of 1406 comparisons (checklists). eBird checklist contributions varied seasonally with lows during winter and summer and highs during migration periods (Supplemental Figure 1). Our analyses included 246 different eBirders who contributed from 1 to 321 checklists.

### Percent error

Across all twenty species, 76 percent of all counts fell short of *R*\* (Figure 1, Supplemental Figure 2), indicating that count data in eBird checklists regularly contained apparent counting errors. eBird checklists with species non-detections excluded (that is, no counts of zero included, even if the species was known to be present that day) had counts below *R*\* values by a median of 29.1% but errors were quite variable across species (Figure 1a), with median absolute deviations of *percent error* averaging 44.6% (Supplemental Table 1). At the extremes, count differences across waterbird species ranged from negative 99% for severe under-counts to more than 3788% too large. In real numbers, counting differences ranged from being too low by 1443 to too high by 1048 (both for Northern Shoveler; Figure 1b). Median percent error was negative, indicative of undercounting, for all waterbird species except the uncommon Surf Scoter (0%; *R*\* was always less than 11).

**Figure 1.**
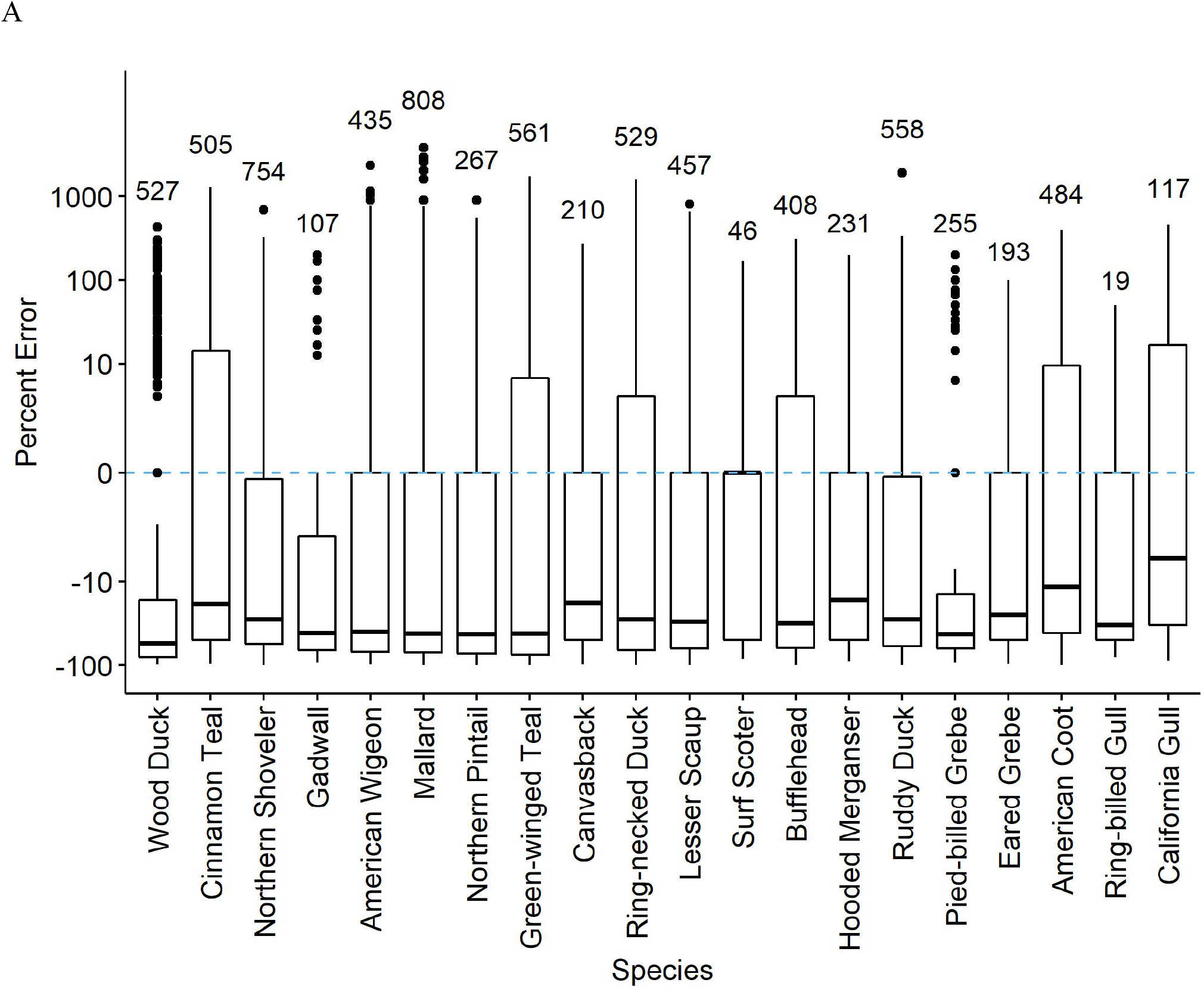
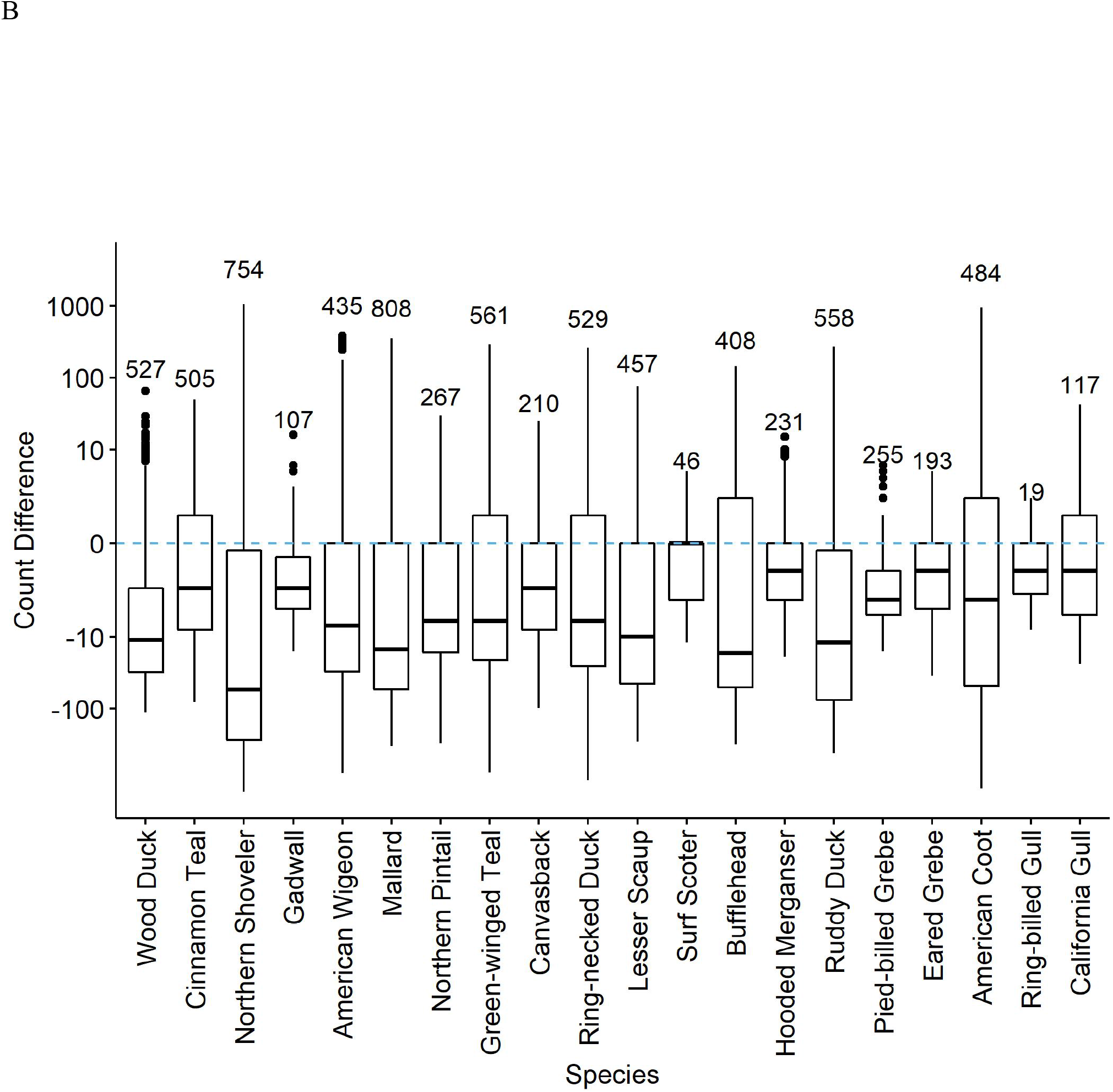
Percent error (A) and count differences (B) in counts of 20 waterbird species reported on eBird checklists at the Philomath Ponds, Oregon USA, 2010-2019. Medians, quantile plots and outliers are indicated, as well as number of checklists reporting counts of each species. Only checklists reporting counts greater than zero were included. For checklists including counts of zero on dates when *R*\* counts were non-zero, see Supplemental Figure 2.

Percent error, when averaged across species and all observers, was fairly consistent at 30% when counts were 30 or greater. Below thirty, counts were more accurate, being closest to zero error when counts were of 8-10 birds (Figure 2A). Percent error was related to the percent richness (our index of observer skill where higher percentages indicated an observer included more of the species known to be present that day on their checklists) in a curvilinear fashion. Checklists including the lowest richness tended to overcount (Figure 2B). Those including 50% of the expected species undercounted by 50% on average, while checklists including 90% or more of the species reported on R* checklists averaged errors of 15% or less in count.

**Figure 2.**
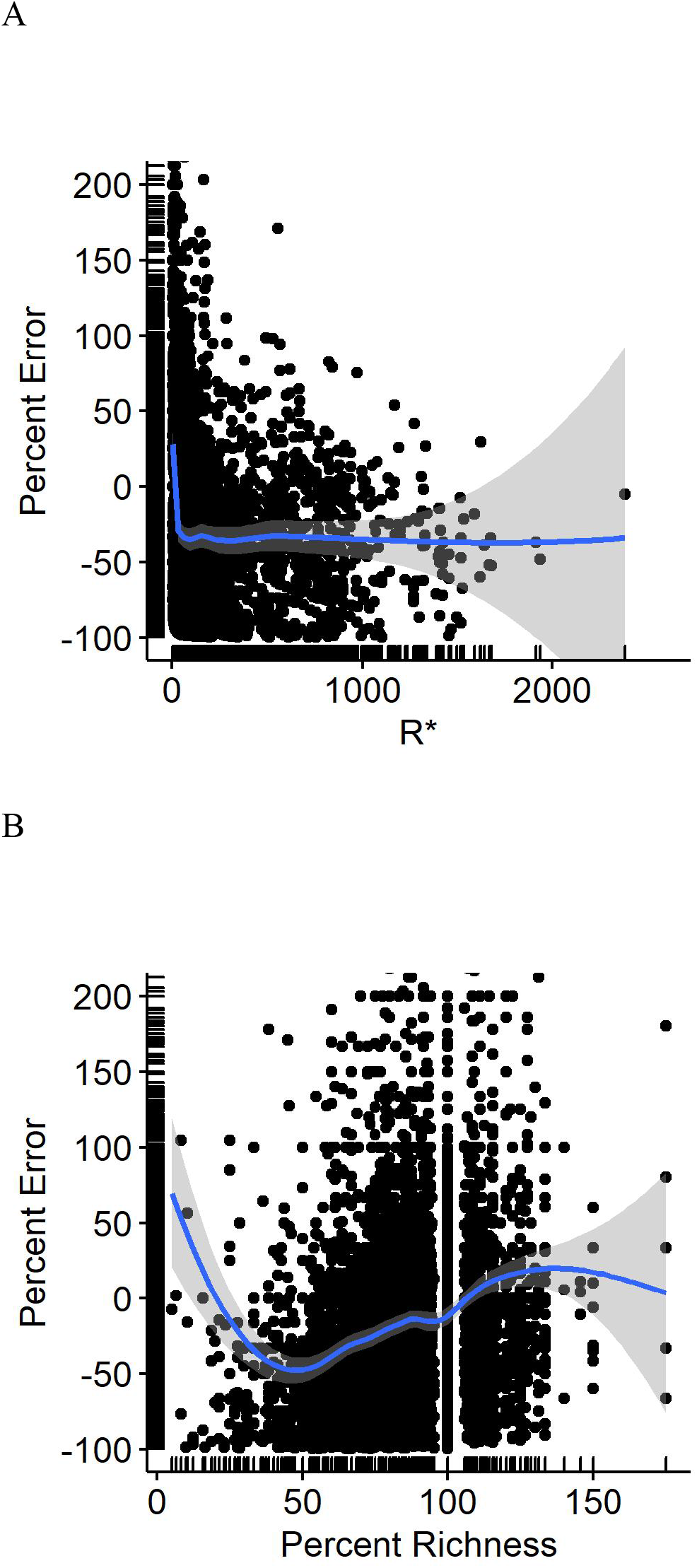
BIC-model predicted percent error in eBird waterbird counts as a function of A) reference (benchmark) counts (*R*\*), and B) percent richness of waterbird species detected at Philomath ponds.

### BIC Top models

In our multi-species mixed-effects model set, our top model garnered 70 percent of the model weight and was over four BIC from the next most competitive model (Table 2). Our BIC top model indicated that a quadratic effect of *R*\* and a linear effect of percent richness best explained variation in percent error.

**Table 2.** Model results of variables most influential on percent error. *R*\*2 is the quadratic of the daily reference (benchmark) count; percent richness is the fraction of the waterbird species present each day that were included on each eBird checklist; duration was the length (minutes) of eBird checklist observation period; starttime was time of day each checklist was initiated; dichromatic was whether each waterbird species exhibited plumage dichromatism or not; date2 was the quadratic of day of year; and proRichness was the total species detected by WDR on each date. See supplemental materials for the full model results.

|  | df | Log likelihood | BIC | delta | weight |
| --- | --- | --- | --- | --- | --- |
| **R\*2_Percent Richness** | 7 | -9751.3 | 19565.0 | 0 | 0.696 |
| **R\*2_Percent Richness_duration** | 8 | -9749.0 | 19569.4 | 4.44 | 0.075 |
| **R\*2_Percent Richness_starttime** | 8 | -9749.2 | 19570.1 | 4.72 | 0.066 |
| **R\*2_Percent Richness_dichromatic** | 8 | -9749.4 | 19570.4 | 5.19 | 0.052 |
| **R\*2_Percent Richness_date2** | 9 | -9745.0 | 19572.7 | 5.41 | 0.047 |
| **R\*2_Percent Richness_proRichness** | 8 | -9750.7 | 19573.0 | 7.79 | 0.014 |

Seasonality in bird numbers was also captured when the second-order *R*\* was included as the most informative variable predicting *percent error*. Numbers of all species varied considerably across each year (Figure 3). Likewise, total waterbird abundance varied several-fold from its nadir in June to a maximum in October and November (Supplemental Figure 3). Yet, total waterbird abundance was rarely an informative variable in our model sets. Only in counts of American Coot did it appear in the most parsimonious models (in combination with percent richness). In California Gull, waterbird abundance appeared as an informative variable but only in a weakly competitive model (19% of the model weight).

**Figure 3.**
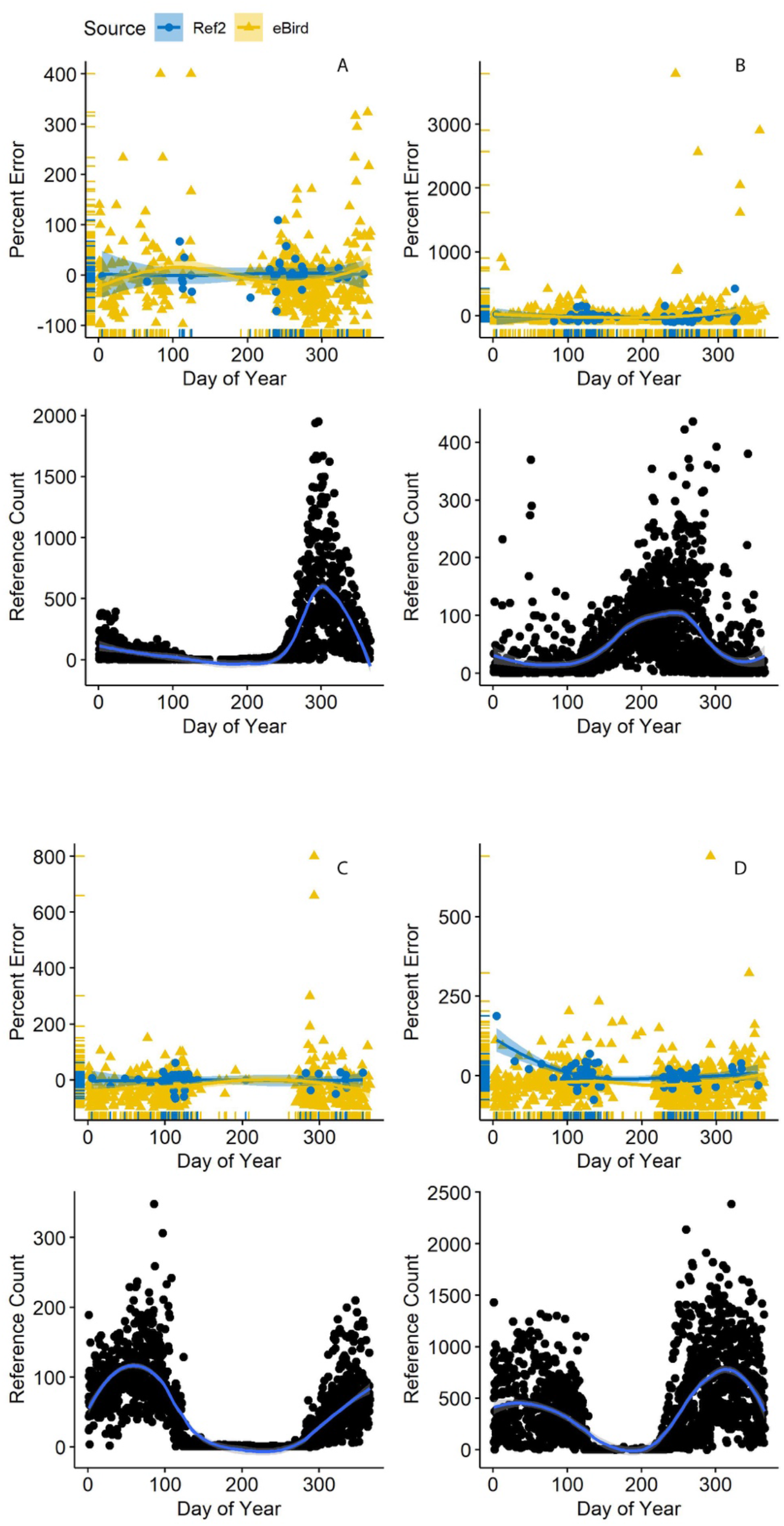
Variation in reference (benchmark) counts (*R*\*) as a function of date (lower panel) and counts reported in eBird (gold triangles in upper panel) alongside second-visit counts (Ref2; blue circles) at Philomath ponds, Oregon USA, 2010-2019. Counts in the upper panels are indicated with respect to the *R*\* count (zero line) each day. Loess regression lines with 95% confidence intervals are included. A) American Coot; B) Mallard; C) Lesser Scaup; D) Northern Shoveler.

Within the species-specific model sets, the combination of *R*\* and percent richness carried most of the model weight (mean=0.83, SD=0.18) in 13 of our 18 non-gull species (Supplemental Table 2). For gulls, top models struggled to outcompete the null. Altogether, R* and/or percent richness were in the top model sets for 17 of 18 non-gull waterbirds.

### Associations with bird characteristics

Within our full model, bird characteristics were rarely influential on percent error (Table 2). Similarly, species-specific models rarely discovered bird traits to be informative variables (Supplemental Table 2).

### Observer effects

Our models often identified percent richness as an influential variable on percent error, so we related percent richness to percent error as means across all checklists contributed by each observer (Figure 4a). The two were positively related, yet only six of the 246 observers averaged *percent errors* of less than 10%. The range in percent error for observers detecting 90% or more of waterbird species was actually greater than the range for observers who detected less than 60% of species, indicating that percent error alone is an unreliable predictor of counting accuracy. The relationship was not necessarily driven by site experience because four of the six observers with the most accurate counts were contributing very few checklists (Figure 4b).

**Figure 4.**
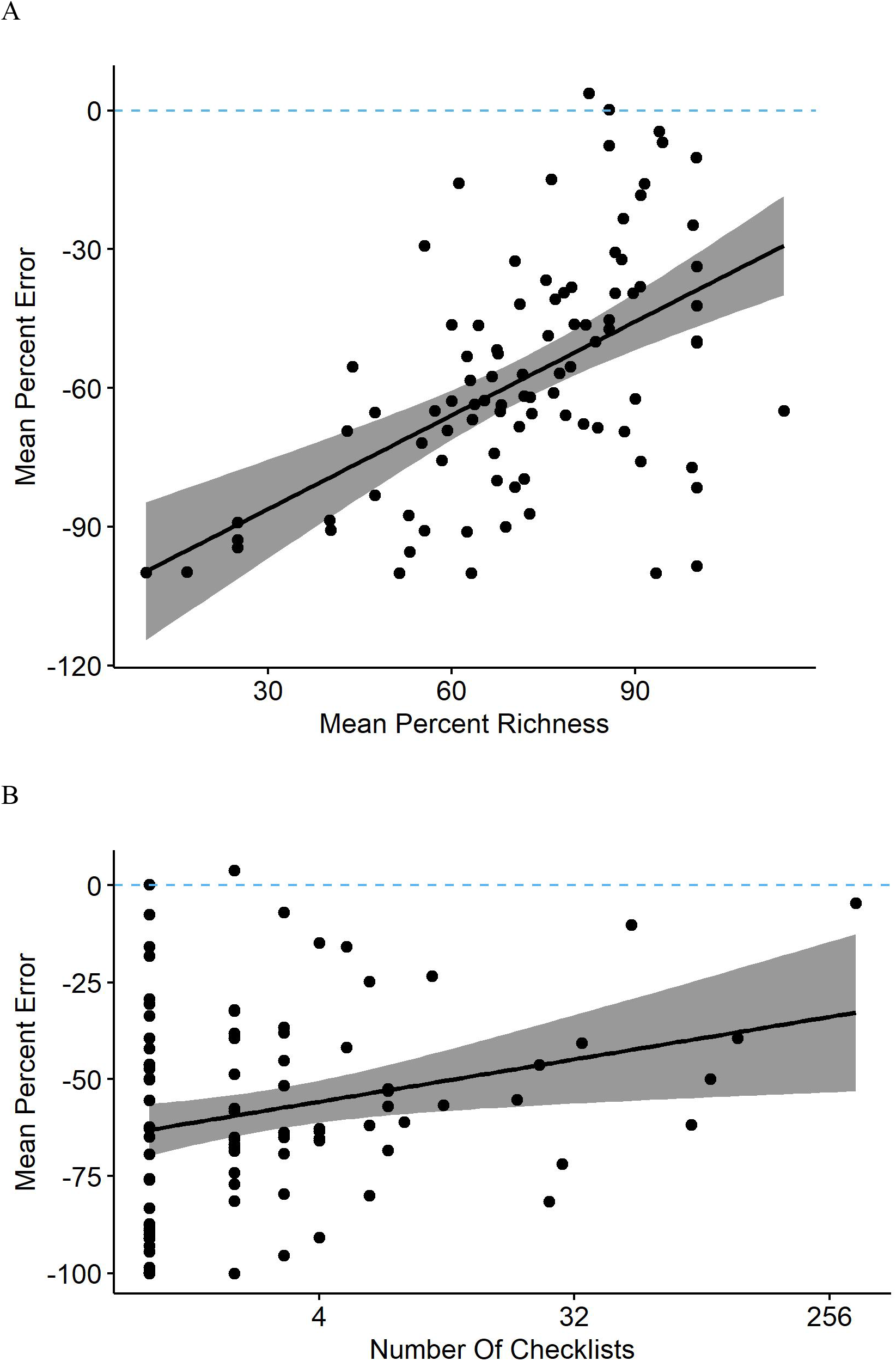
A) Observers reporting a greater percentage of waterbird species present at Philomath ponds, Oregon USA, tended to have lower percent counting errors in their eBird checklists (linear regression and 95% confidence intervals; y=−110 + 0.68x). B) Observers submitting more total checklists tended to have lower counting errors (y=−60 + 0.17x). Note that these are means of all applicable checklists for each observer, so each point represents a unique observer.

We then selected checklists from the ten observers who contributed the most. Those checklists also showed evidence of undercounting. In nearly all 20 species, percent error was 10 to 60% greater than even the Ref2 counts (Figure 5). Percent error was highly variable across species. In some species, such as American Coot, three of the 10 observers reported counts averaging very near the Ref2 counts, whereas in others, such as Pied-billed Grebe, all observers undercounted by at least an average of 20%. Again, percent error was highly variable in all species even when median percent error did not deviate far from zero.

**Figure 5.**
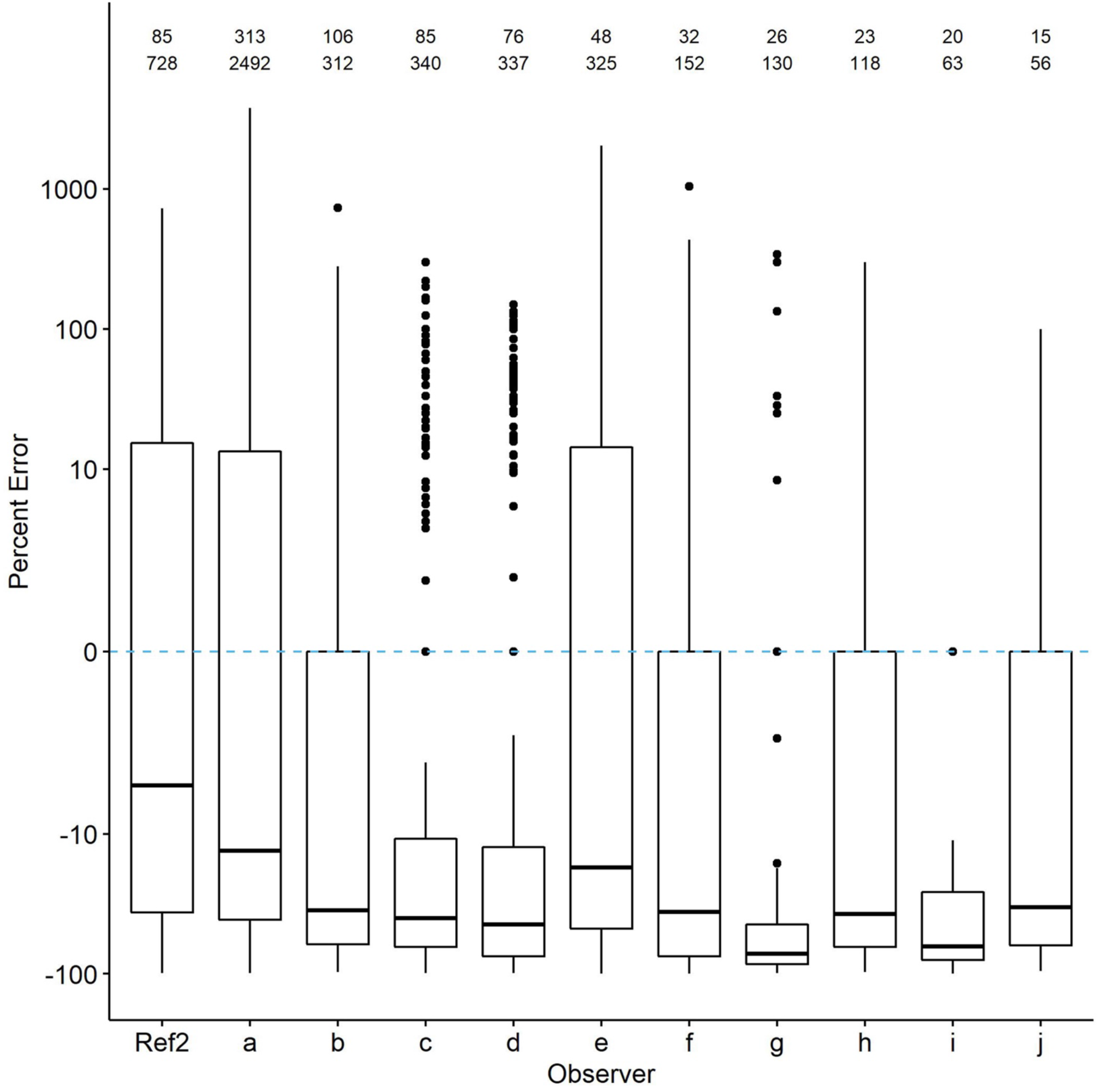
Comparison of percent count errors in eBird checklists contributed by the 10 observers with the most checklists (top row of numbers) and waterbird observations (second row of numbers; each checklist includes multiple species). The zero line is *R*\*. Ref2 is the second-visit data from WDR. Quantile plots show the median, 25^th^ percentiles as boxes and whiskers, plus outliers. Species-specific plots are available from the authors upon request.

### Community visualization

We visualized characterization of the richness and abundance of the daily waterbird community with NMDS through ordination of checklists (grouped by observer) in species space. Observers characterizing the community and its species abundance patterns similarly to *R*\* fell nearer to *R*\* whereas those positioned increasingly further from *R*\* described the community in increasingly dissimilar details. In both January (Figure 6a) and October (Figure 6b) high inter-observer variability in how their checklists characterized the waterbird community led to a general lack of clustering near *R*\*. In both months, observers reporting more species, contributing more checklists, and surveying for more time tended to group nearer *R*\*. The collective average, 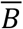, was nearer *R*\* than any individual observer during January but one observer was closely positioned near 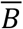 during October. Removal of checklists from the observer contributing the most data had minimal effects on results.

**Figure 6.**
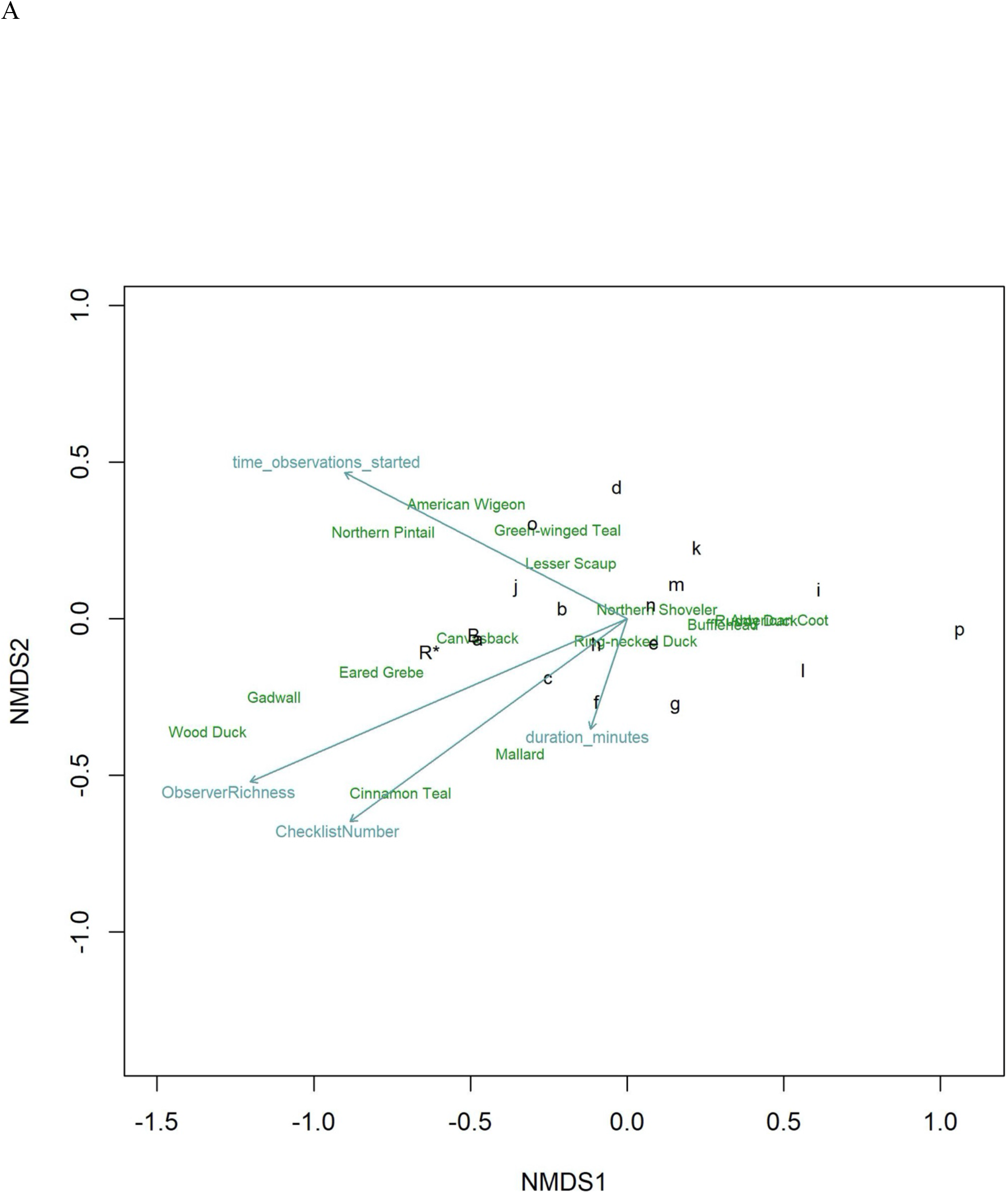
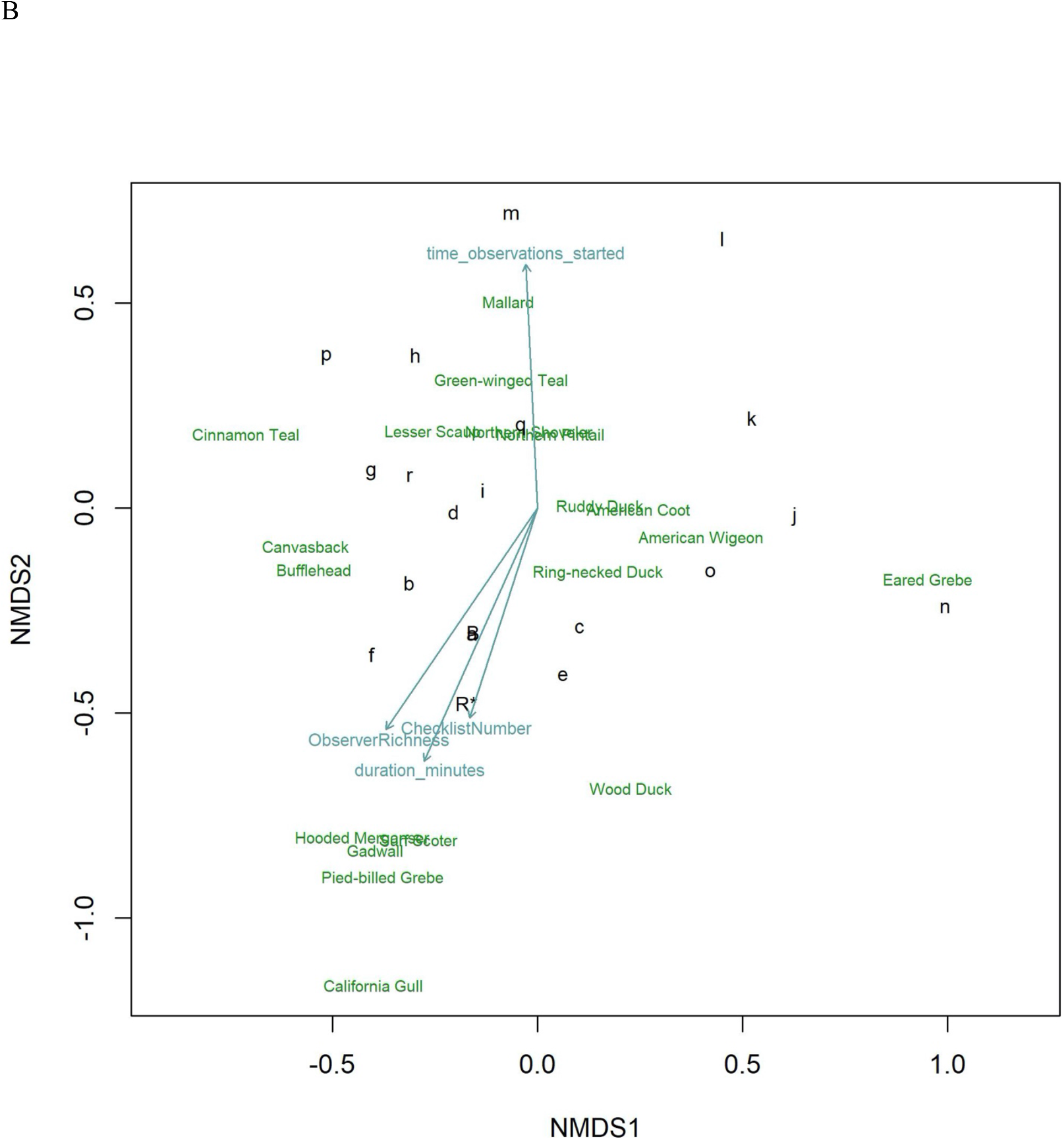
Ordinations using NMDS of eBird checklists characterization of the waterbird community during A) January and B) October at Philomath ponds, Oregon USA, 2010-2019. The most influential vectors included Observer Richness (percent of known richness reported on each checklist), Checklist Number (total number of checklists per observer), observation start time each day, and the duration of each observation period. Relative positions of species in species space are noted by species English names. Benchmark counts are noted by *R*\*. Individual observers are noted by lower case letters; those nearest to *R*\* produced characterizations of the waterbird community most like *R*\*. 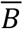 is the collective average of eBird checklists, showing that from the perspective of generally characterizing the community, averaging across checklists contributed by many observers aligns more closely with *R*\* than do checklists from most individual observers, although observer *a* occupies nearly the same location in species space.

## Discussion

Benchmark data are often designed to understand temporal change in biodiversity (Curtis and Robinson, 2015; Curtis et al., 2016; Robinson and Curtis, 2020). Here, we show that they can also be used to establish standards that aid in quantification of potential errors in citizen-science data. Through comparisons with such a standard, we discovered that bird count data contributed to eBird from our study site were consistently biased toward undercounting. Counts averaged approximately 30% too low whenever benchmark counts were of 30 or more birds. Importantly, however, errors exhibited high variability across species and observers. Benchmark data like ours can subsequently inform decisions regarding what subsets of data should be selected to most rigorously address particular scientific questions or management decisions, analogous to how checklist calibration indices help researchers choose suitable eBird checklists based on site- and time-specific expectations of species richness (Yu et al., 2010; Kelling et al., 2015; Johnston et al., 2018). Yet, situations in which such informative standards may be developed and compared appear to be rare currently.

Our study site presented a unique opportunity to compare bird count data contributed to a citizen science database (eBird) with benchmark reference data collected by a professional observer focused on generating accurate daily counts. Characteristics of the site, where all birds were in the open and identified by sight, minimized issues of availability and therefore the need for detectability adjustments to compare counts. Data were contributed by 246 observers and included 676 dates across 10 years, providing an unusual opportunity to explore patterns and potential sources of error. Although the extent to which our results may be generalized to other sites remains unclear given the rarity of opportunities like this one, the situation probably represents a best-case scenario given that birds were in the open and easy to observe. Despite the advantages, count differences in 20 species of waterbird were highly variable across the calendar year, species, and observer. Coefficients of variation were high, averaging 6.6 across the 20 species and ranging from 1 to 35.6. For comparison, in an experimental study of observer counting errors of singing birds, which should have been much harder to detect and identify but had a lower range of abundances than our waterbird community, coefficients of variation averaged 0.1 (Bart, 1985).

Our quantification of counting error is actually conservative because we excluded counts of zero on eBird checklists, even for species known to be present. We did so to minimize the potential confound of misidentifications and reporting errors (failing to enter a count for a species that was actually observed) from our analysis of counting errors. Yet, it is possible that some fraction of 100% undercounts were indeed counting errors in the sense that the species was one that observers were knowledgeable enough to identify but failed to count or report. The median percent error across the 20 species was −48.6 plus or minus 50.9% (MAD) when zero counts were included versus −29.1 plus or minus 44.6% when zero counts were excluded. Inclusion of zero counts, therefore, has a large influence on the median, but percent errors were highly variable regardless.

Our top overall mixed-effects model carried nearly 70% of the model weight and contained only two variables. The species-specific *R*\* count as a quadratic, which captured the seasonality in numbers present at the site, was the most informative variable when combined with a linear effect of percent richness. The inclusion of *R*\* indicates that eBird count data were related to the benchmark numbers but that other factors were also influential. Checklists with a more complete list of the species known to be present each day had lower counting errors. Yet, checklists including 100% of expected species still undercounted by an average of 15%. Count differences on checklists from the ten observers who most often visited the site were still exhibiting undercounts even compared to the Ref2 values, which were benchmark counts made later each day during weeks with high levels of migratory movements.

We documented strong directional bias toward undercounts and also a smaller percentage of large overcounts, leading to inconsistent patterns in count differences across species. Our comparisons revealed that undercounting was pervasive, yet very large numbers of a species being present sometimes led to severe overcounting as well. Interestingly, the influence of number of birds appeared to be species-specific. The total number of waterbirds of all species present on a given day was not an influential variable in our overall model explaining percent error, except for one species, American Coot. This pattern suggests that count differences were unlikely to have been caused by observers being overwhelmed by the total number of birds to observe, identify and count. Instead, it appears that each species presented different challenges to observers. Given that our models rarely identified species’ traits as being informative, it remains unclear what species-specific factors are responsible.

The degree of variability across species in count differences should influence potential decisions regarding use of eBird count data. Our analyses clearly reveal that off-the-shelf acceptance of count data for assessments of absolute abundance should be done with great care and thoughtfulness. In addition, if researchers wish to avoid focus on absolute abundance by instead evaluating relative abundance, our results suggest further caution is warranted. We found great interspecific variability in count differences. That is, although bias was nearly uniformly directional toward undercounting, the magnitude of undercounts varied substantially across species indicating that processes generating errors are inequivalent across species. Therefore, judging differences in one species’ abundance relative to others requires careful thought. If explorations of relative abundance are focused on within-species changes across sites, care is also warranted because we found substantial differences among observers in count accuracy. If different sites have different observers, then error/bias processes will be expected to be different as well. Effective use of relative abundance data depends on assumptions of consistent errors across species and sites, which appears to be largely untrue in our data. Further exploration of techniques to determine the degree to which assumptions of similar counting errors across species might be relaxed to preserve the utility of relative abundance analyses are warranted. The use of abundance categories could be explored to maximize the information content gleaned from count data.

What role might species misidentifications have played in counting errors? Count differences were regularly so large that we conclude species misidentification was unlikely to be an important factor. Probably the most challenging identifications involved female or eclipse-plumaged ducks, which observers might ignore and exclude from checklists if identification is uncertain. We consider such omissions to be unlikely for at least three reasons. First, degree of dichromatism was uninformative in our models explaining percent error. Second, assuming that females represent approximately half of each species present during most months of a year, count differences might be expected to average 50% if males were counted accurately but females were not. Instead, percent error varied widely across species. Finally, count differences of monochromatic versus dichromatic species were not obviously different. However, it is possible that observers were more accurate for some species than others because of paying greater attention to unusual or favorite species (Schuetz and Johnston, 2019). At our site, most charismatic species of great interest to birders are rarities and so were not included in our analyses. Counts of Surf Scoter, a species that occurs during a narrow window of time in fall, were generally accurate, but we cannot attribute the accuracy to celebrity alone given its occurrence in such small numbers.

Based on our analyses of count differences at this site, it appears that count data on eBird checklists from similar situations should be used with great care and thoughtfulness. Aside from a predominantly directional bias toward undercounts, we found few consistent species-specific patterns in percent error. Errors differed in magnitude across species, observers, and time of year. Therefore, development of some type of calibration effort, where checklist numbers are adjusted to more closely approximate species-specific abundances poses an interesting challenge. The variability in raw count data suggests that tracking trends across time without additional steps to filter data or analytically adjust for noise could be especially problematic. Depending on the particular scientific question of interest, needs for precision might decline, so other analytic approaches could be implemented. For example, if abundances can be binned into categories and approaches such as ordinal or quantile regression used (Ananth and Kleinbaum, 1997; Koenker and Hallock, 2001; Howard et al., 2014), less precisely defined trends over time might be identified. Furthermore, our observation that percent richness, which we assume to be a correlate of observer experience, was often an informative variable, suggests that additional exploration of count calibration approaches for data contributed by the most experienced observers might be informative.

If questions about patterns in abundances among species in the waterbird community are of interest, our NMDS ordination results suggest that combining checklists across multiple observers may produce results closer to those generated by professional benchmark data. The vectors in NMDS results may also inform decisions about which criteria to use when filtering data to maximize inclusion of checklists with the greatest value for specific scientific questions. For example, the waterbird community at our site was better characterized by observers who included more species on their checklists, invested more time searching the site each time, and contributed more checklists overall. Although species-specific numbers remained inconsistently related to the *R*\* counts, the level of general characterization of the entire community was improved. In a detailed comparison of eBird data with structured survey results near Sydney, Australia, overall characterization of the bird communities was similar as well, but the collectively greater effort expended by eBirders resulted in discovery of a greater number of uncommon species (Callaghan et al., 2018).

Determining the extent to which results from our site and observers may be generalized more widely will require identification of other sites with benchmark data sets. We also recommend further investigation of approaches for identifying checklists with higher probability of having the most accurate count data. New approaches for categorizing checklists based on expected numbers of species have recently been developed but it remains unclear if these same criteria also apply to bird counting accuracy (Callaghan et al., 2018). Our index of checklist quality was based solely on the percent of species reported on checklists that were also detected that day by the professional observer. Percent richness was regularly in top models, so does have explanatory influence on count differences. Yet, direct comparisons of data from those observers and the *R*\* and Ref2 numbers still showed substantial differences, primarily of undercounting.

If a sufficiently detailed benchmark data set is available, however, adjustments for seasonal fluctuations in numbers of each species could conceivably be implemented. Such calibrations might be conducted more effectively if individual observers exhibited consistency in counting errors, an issue we have not explored here. It is unknown if observers improve their counting skills over time in the same way that observers are expected to improve abilities to detect species or if temporal stochasticity drives counting errors. A goal could be to develop a count calibration metric for each observer so that it can be extended and applied to counts from sites lacking data from a professional observer if those sites are likely to have similar species composition and relative abundances. However, given the high level of variability in count data we quantified across observers, species and time, such calibration metrics may be quite challenging to develop. Complex models such as the Bayesian hierarchical models using Markov chain Monte Carlo approaches like those implemented with Christmas Bird Count data (Link et al., 2006), might be helpful in the absence of additional information on checklist accuracy and reliability. Our community ordination results suggested that combining data across multiple checklists from multiple observers (the group collective effort) might more closely approximate the community characterization than most single contributors did. Further exploration of similar approaches and sensitivities to checklist characteristics could identify necessary checklist quality criteria that must be met prior to use in such analyses. In the end, use of any checklist count data will be influenced strongly by each project’s specific objectives (Isaac and Pocock, 2015).

We hypothesize that the high variability in species count information on eBird checklists could be influenced by common aspects of birder behavior. Prior to the advent of eBird, most birders, in North America at least, focused their efforts on listing species and watching behavior (Eubanks, Jr. et al., 2004). Intentional counting was done by a small percentage of particularly avid observers, while most others only counted during organized activities such as Christmas Bird Counts (Boxall and McFarlane, 1993). A much smaller percentage contributed count data to scientific projects with structured protocols such as the North American Breeding Bird Survey. eBird has revolutionized the degree of attention birders pay to numbers of birds around them (Wood et al., 2011). It has pushed birders to value data beyond the day’s species list. The novelty of this effort to count all birds every time one goes birding, may contribute to the variability in quality of the count data. Contributors are largely untrained about best practices for counting, especially when birds are present in large numbers, flying, or inconspicuous because they are secretive or available only by sound. We encourage development of additional training opportunities for eBird contributors that improve their knowledge of the value of accurate count data as well as their counting skills. Training improves data quality even for professional observers (Kepler and Scott, 1981).

An indication on checklists in the eBird database that such training had been accomplished might facilitate selection of checklists by researchers who wish to use count data only from trained observers. Furthermore, the addition of a qualitative categorization of counting accuracy for each checklist, designated by the observer at time of checklist submission to eBird, might be useful. Currently, users may code species using presence-absence information instead of counts or select a checklist protocol (incidental) indicating that not all species detected were included the list. A count accuracy designation could allow observers to rate their own level of confidence in the accuracy of their counts or the level of attention they paid to counting accurately, which could serve as additional criteria by which researchers might choose checklists for their particular scientific question. Given that many contributors may not necessarily participate to contribute data useful for abundance analyses but have a variety of other motivations (Boakes et al., 2016), allowing observers to categorize quickly and easily their personal confidence in their count data would be useful.

Finally, exploration of the sources of variation in count data needs additional attention (Dickinson et al., 2010). The potential value of the vast quantities of information from citizen science databases is great. Such data have the potential to be effective at informing conservation and management decisions (McKinley et al., 2017; Young et al., 2019), but a thorough understanding of sources of error should be a priority before their use (Lewandowski and Specht, 2015). An additional strategy that may contribute to refinement of information on count data quality in citizen science databases could be development of a network of sites with trained counters. These marquis sites could be chosen to represent major habitat types where citizen science data are often gathered or where researchers specifically need high-quality information. Creating a network of high-quality benchmark sites would have the added advantage of leaving a legacy of more reliable abundance data for future generations, especially if complete metadata are also preserved.

## Acknowledgements

WDR and TAH were supported by the Bob and Phyllis Mace Professorship and the College of Agricultural Sciences. RAH was supported by NSF Grant 1910118. We thank the many birders who contributed their data to eBird and the many scientists who created, maintain and continue to improve eBird. Dennis Lewis and Philomath Public Works permitted access to the site, not only to us but to more than 250 birders. We thank our Reconfiguration Grant Group for helpful discussions.

---

Supplemental Table 1. Species-specific measurements of central tendency and variation in percent counting errors. A) excluding species non-detections from checklists; B) including species non-detections (zero counts) in checklists.

Supplemental Table 2. Species-specific BIC model results. Full model results are presented for each species alphabetically.

Supplemental Table 3. Full mixed-effects model results supplementing the abbreviated results presented in Table 2.

## Supplemental Figures

**Supplemental Figure 1.**
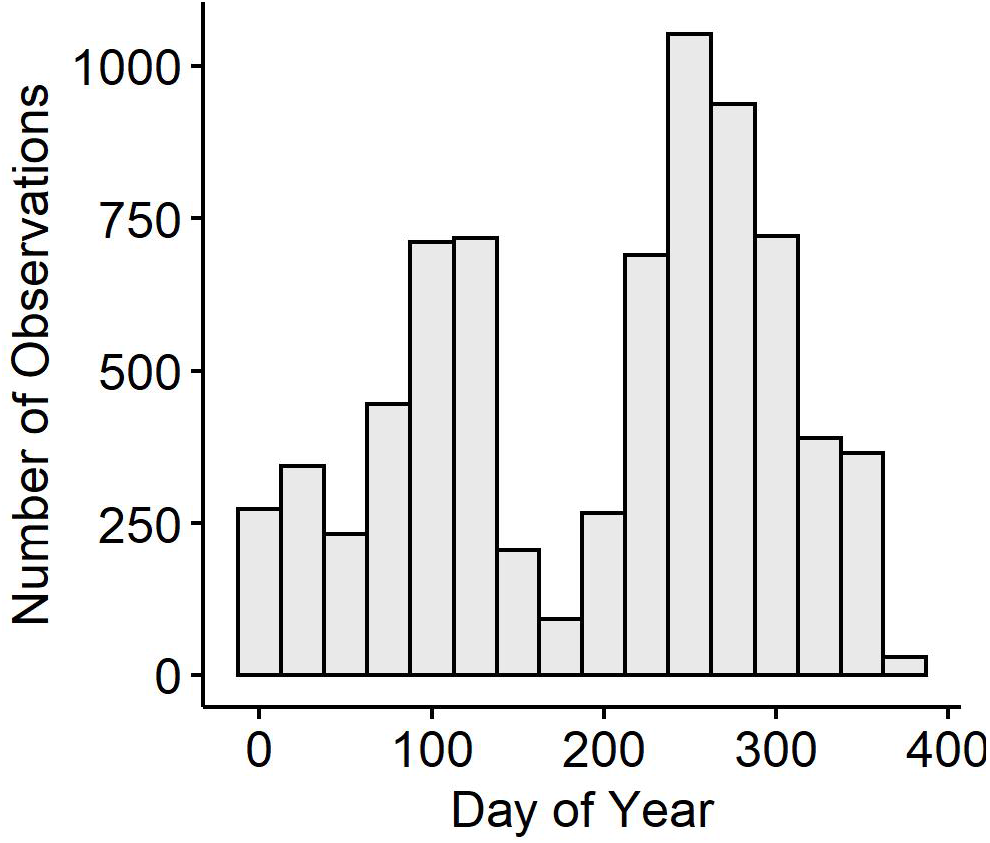
Number of eBird checklists contributed for the study site at Philomath Ponds, Oregon USA, 2010-2019, as a function of day of year.

**Supplemental Figure 2.**
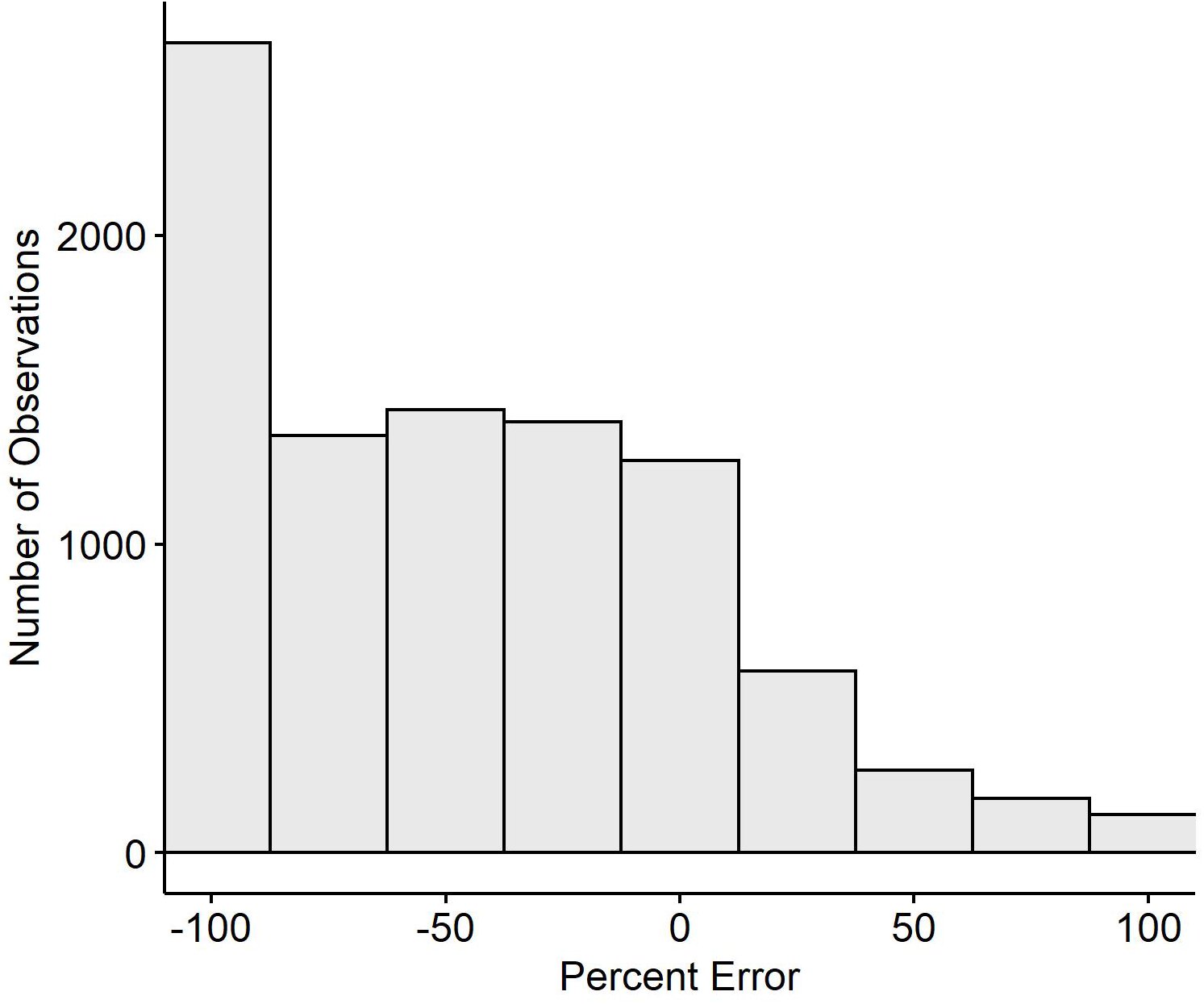
Counts of waterbirds in eBird checklists included in our analyses as a function of their percent error.

**Supplemental Figure 3.**
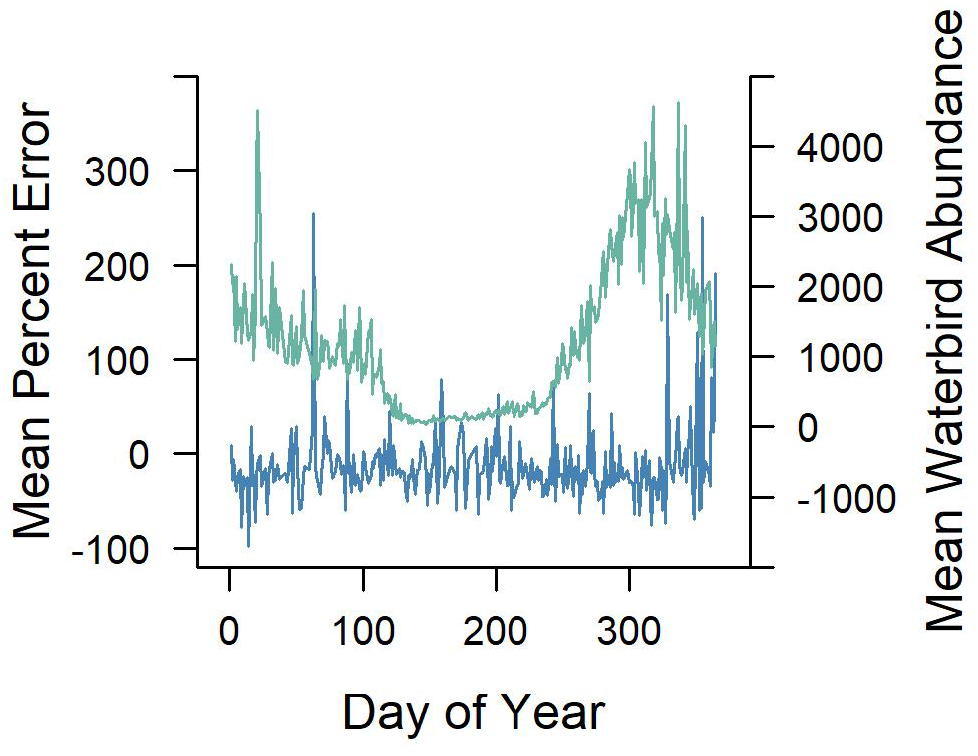
Relationship between mean percent error on eBird checklists (blue line) and mean waterbird abundance (green line) as a function of day of year at Philomath ponds, Oregon USA, 2010-2019. Waterbird abundance is the mean of all the counts (*R*\*) of all of the possible 20 study species present each day across the 10 years.

