## Supplementary figures and images for "Duck, duck, goose: Benchmark bird surveys help quantify counting errors and bias in a citizen-science database"

### AmericanWigeon_Julian_PercentError.jpg

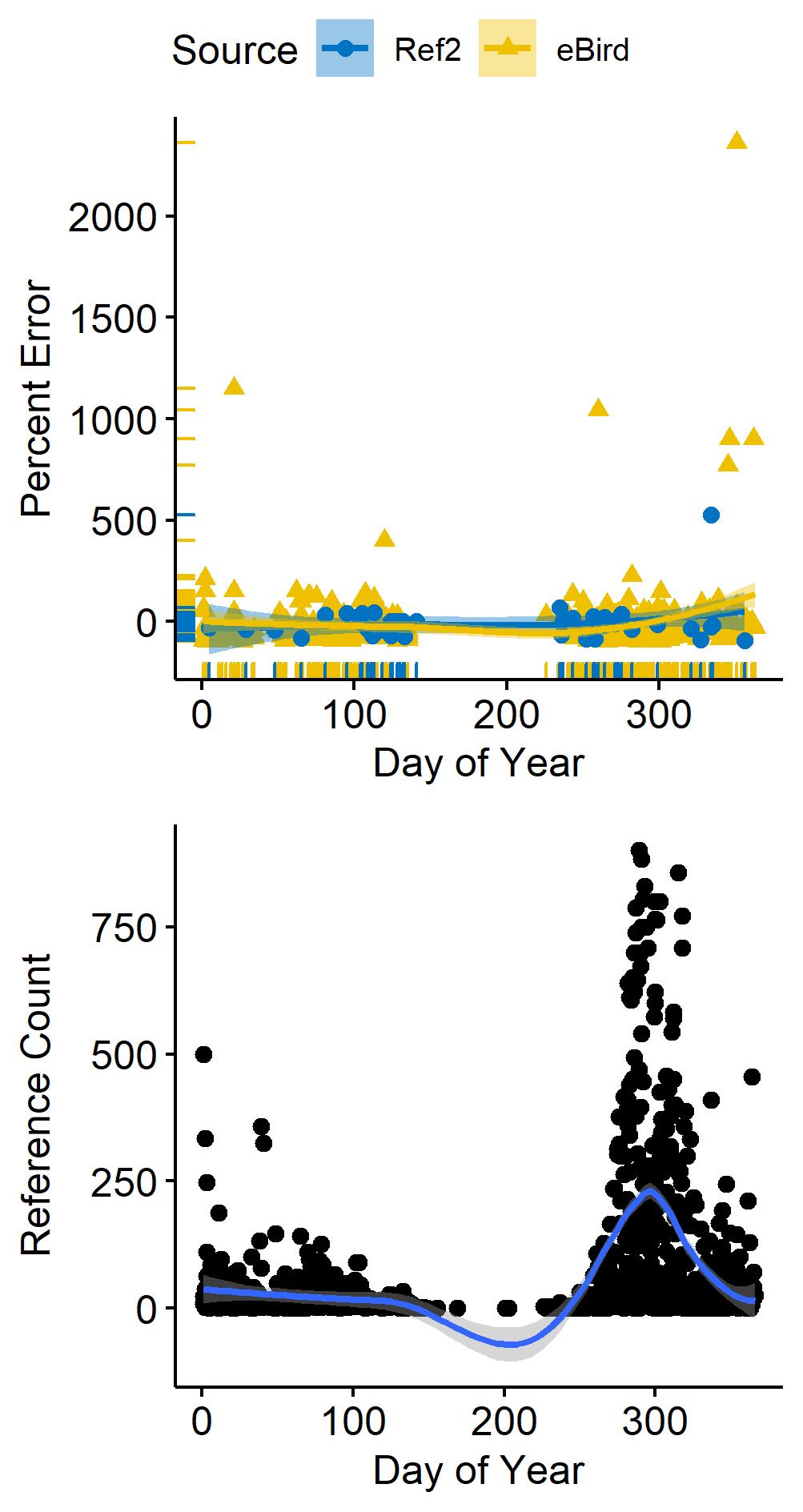

### Bufflehead_Julian_PercentError.jpg

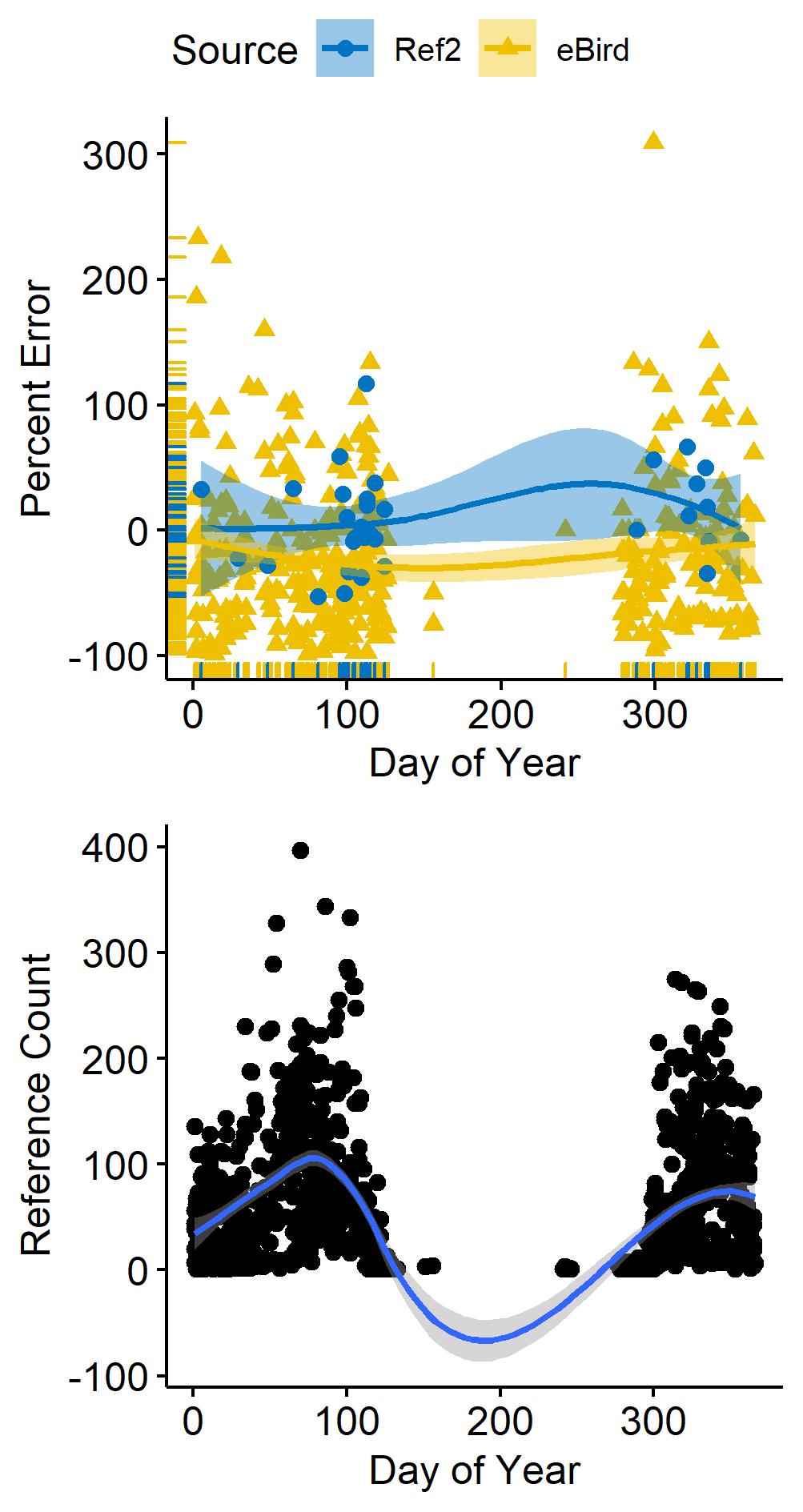

### CaliforniaGull_Julian_PercentError.jpg

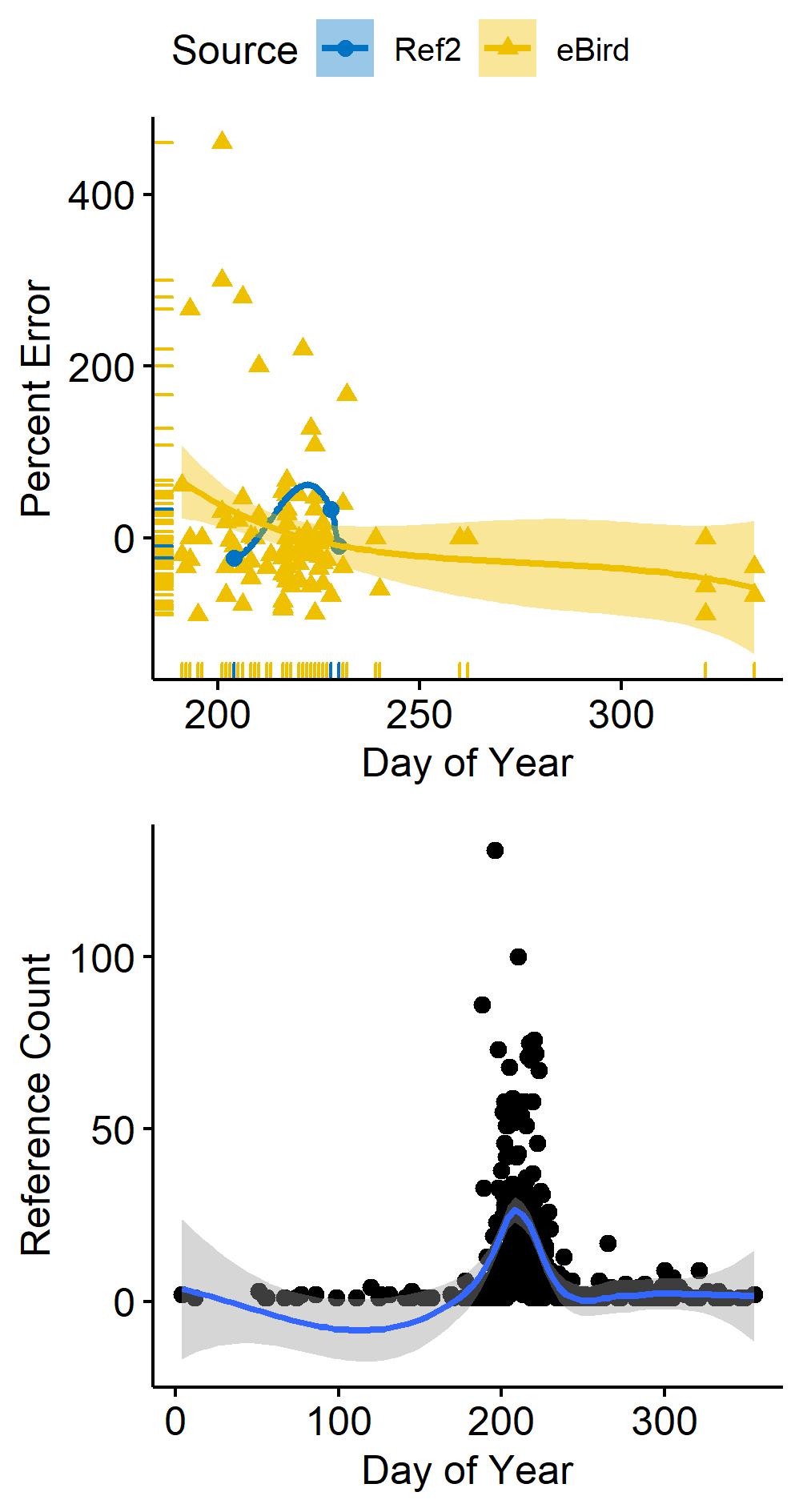

### Canvasback_Julian_PercentError.jpg

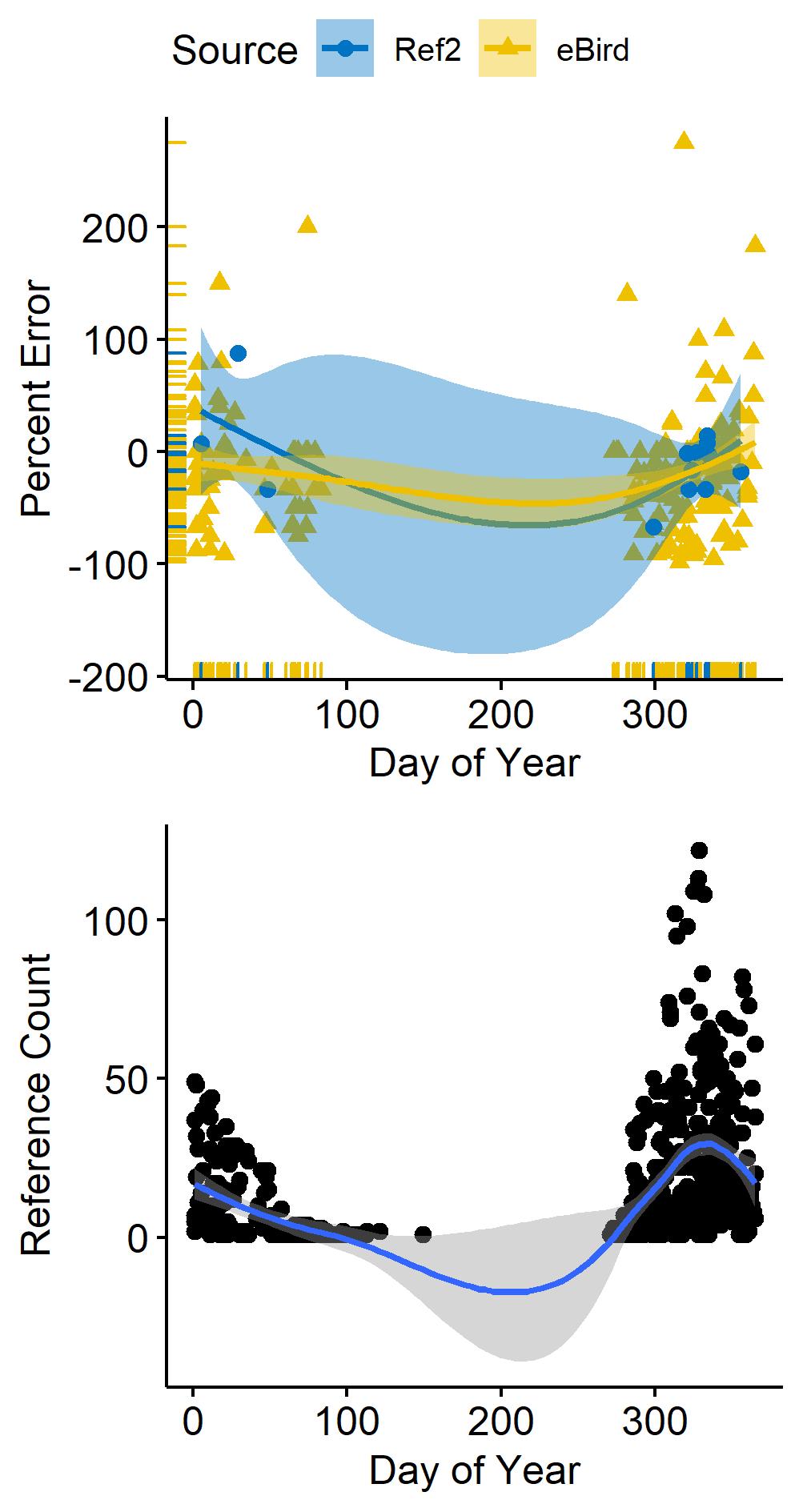

### CinnamonTeal_Julian_PercentError.jpg

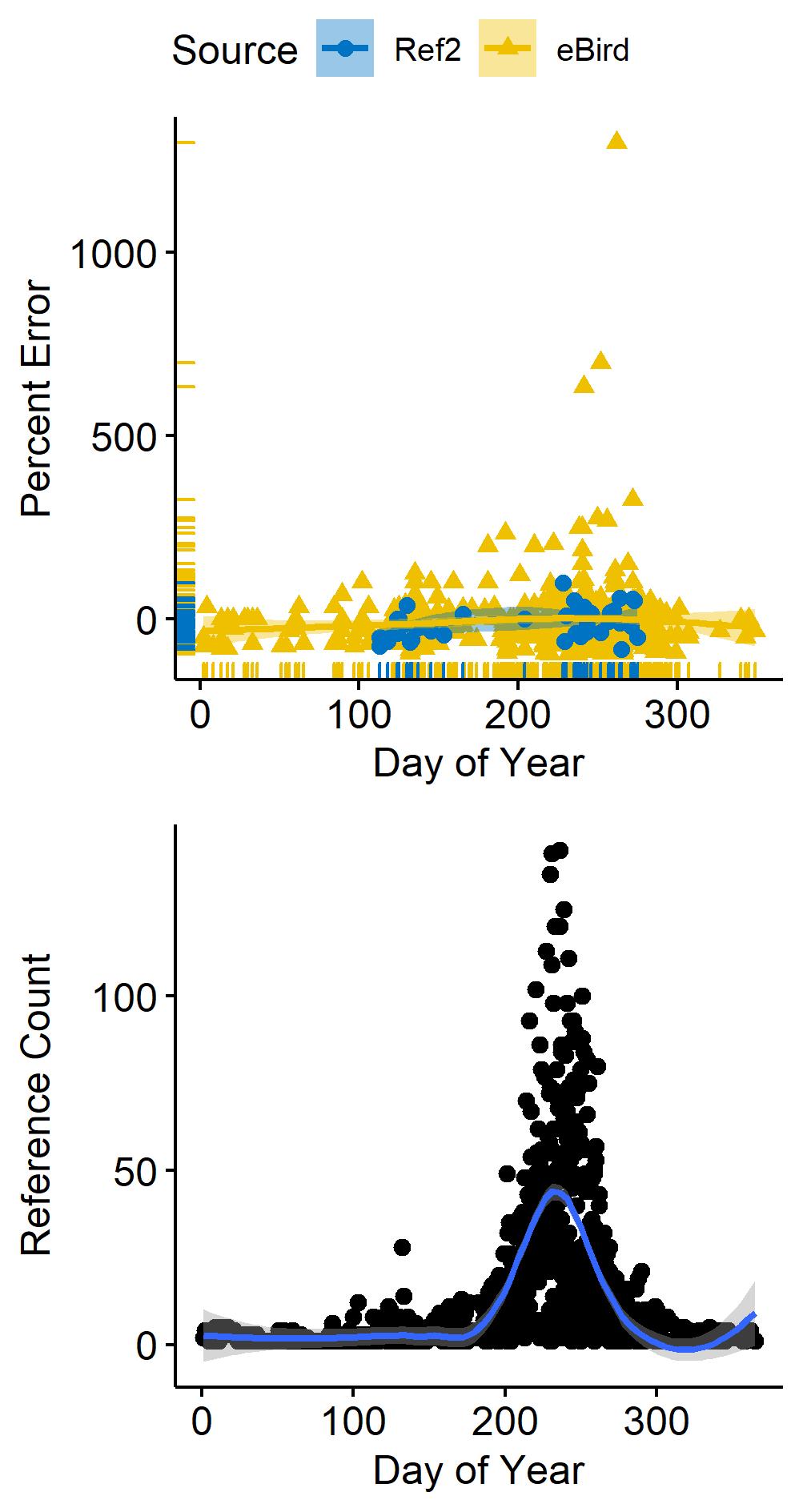

### EaredGrebe_Julian_PercentError.jpg

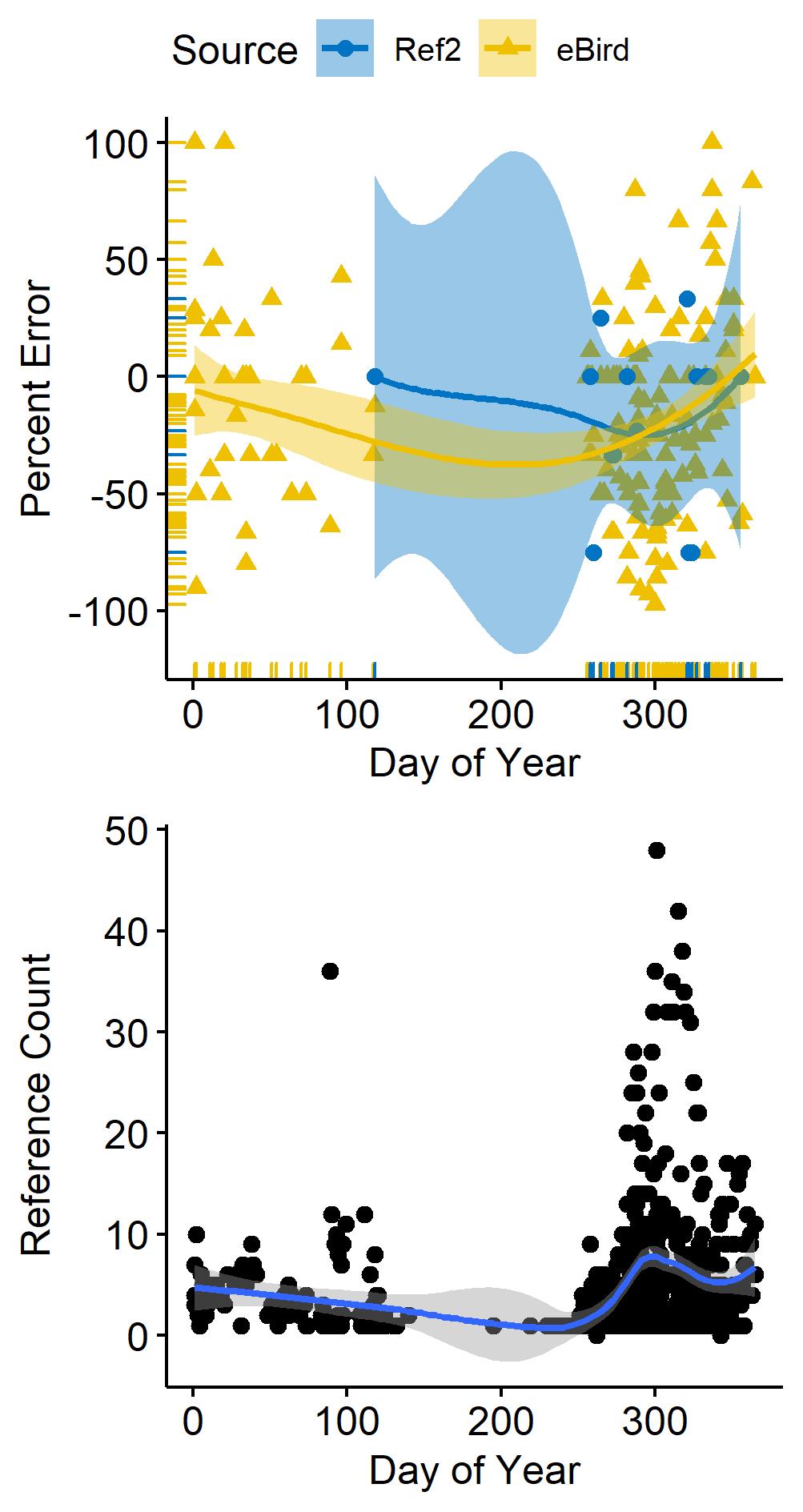

### Gadwall_Julian_PercentError.jpg

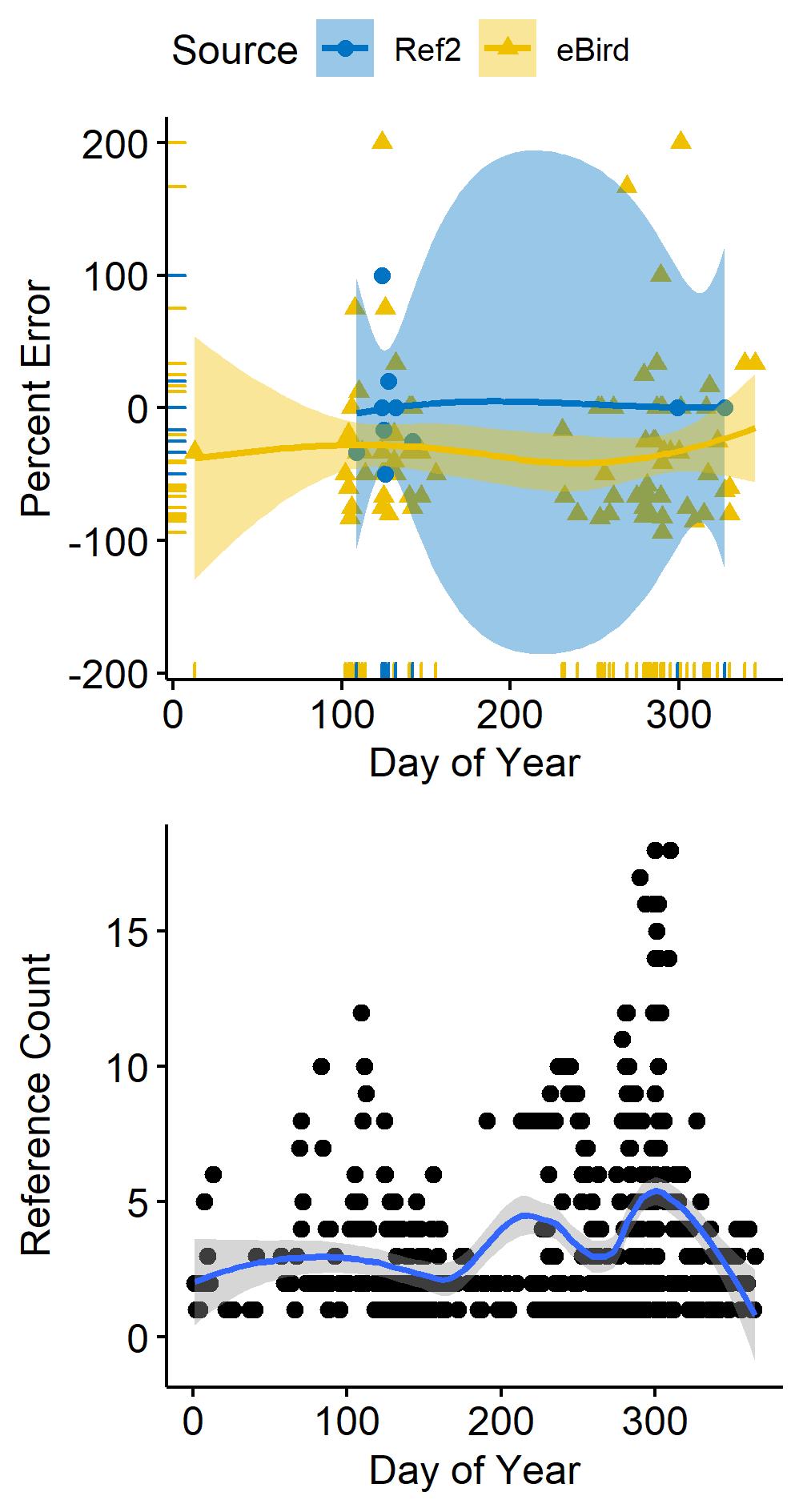

### GreenwingedTeal_Julian_PercentError.jpg

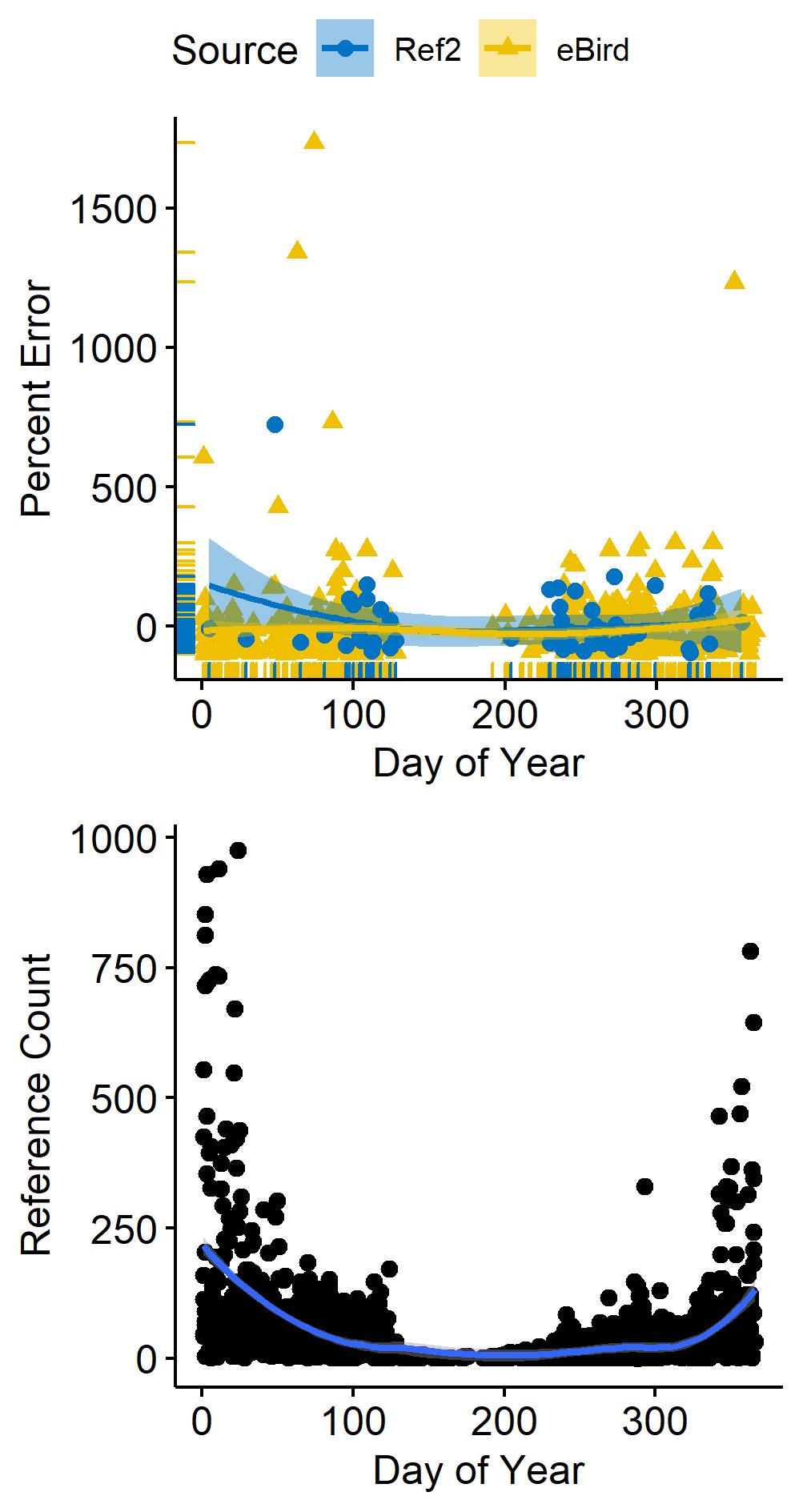

### HoodedMerganser_Julian_PercentError.jpg

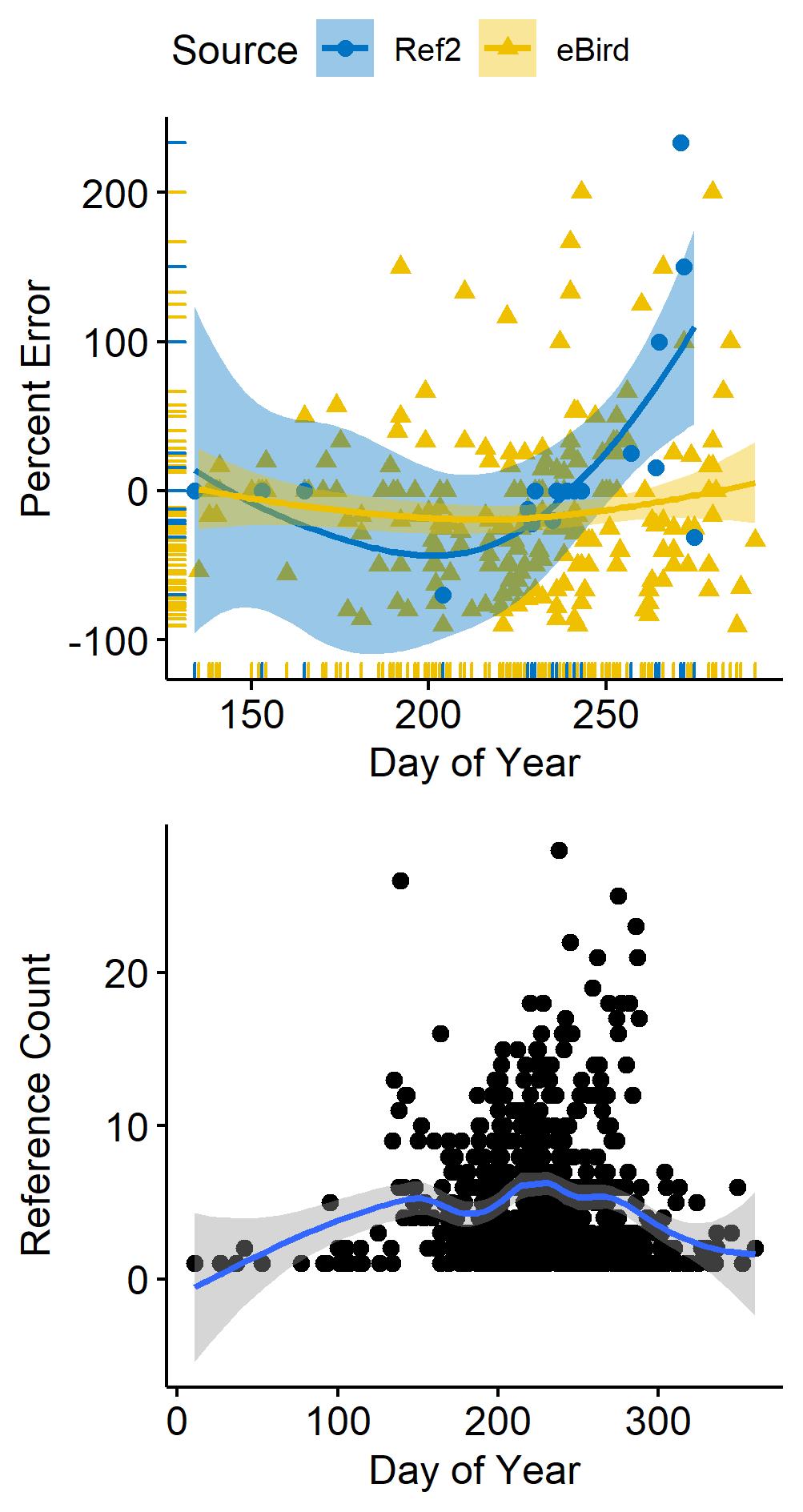

### NorthernPintail_Julian_PercentError.jpg

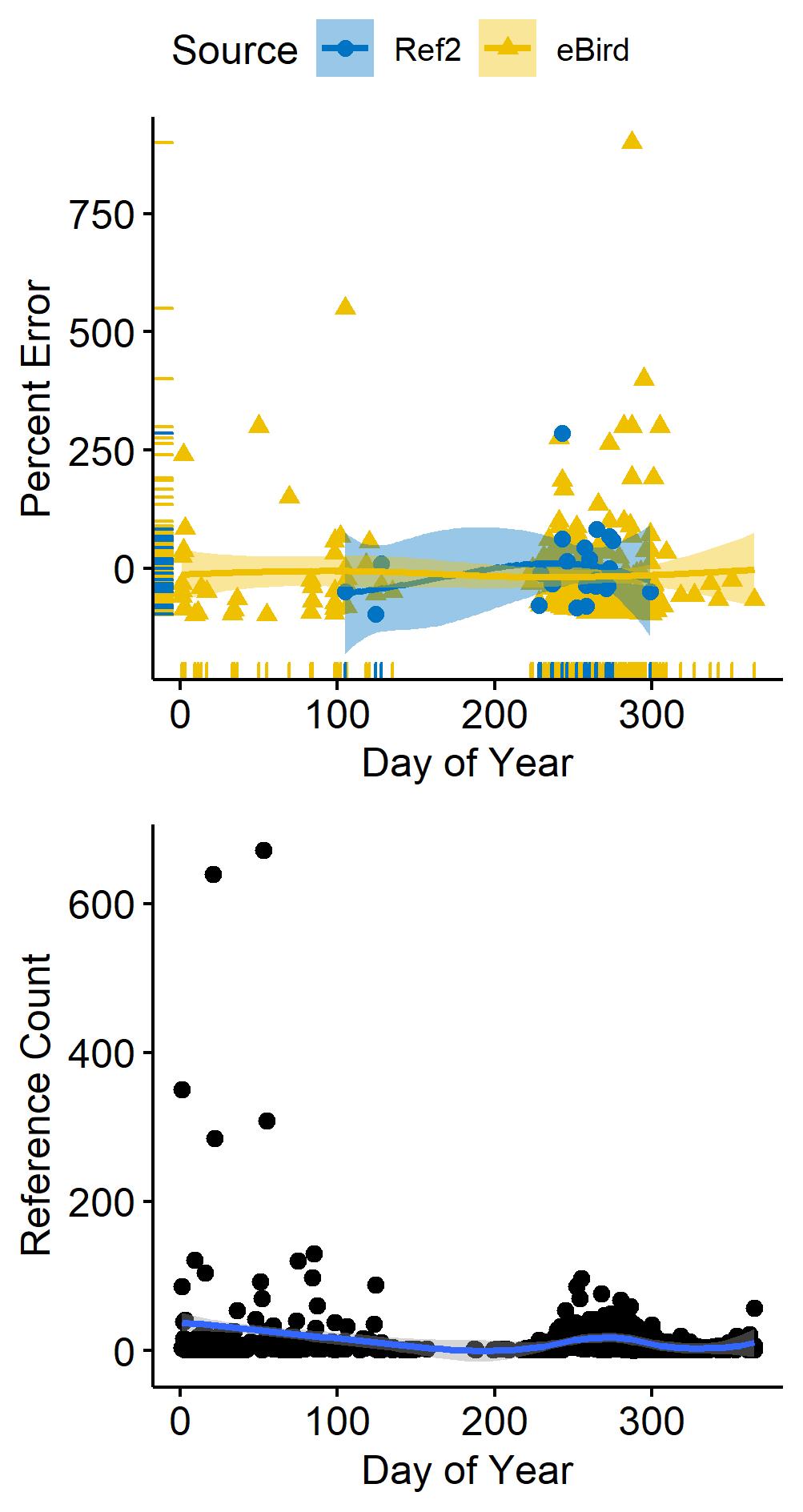

### PiedbilledGrebe_Julian_PercentError.jpg

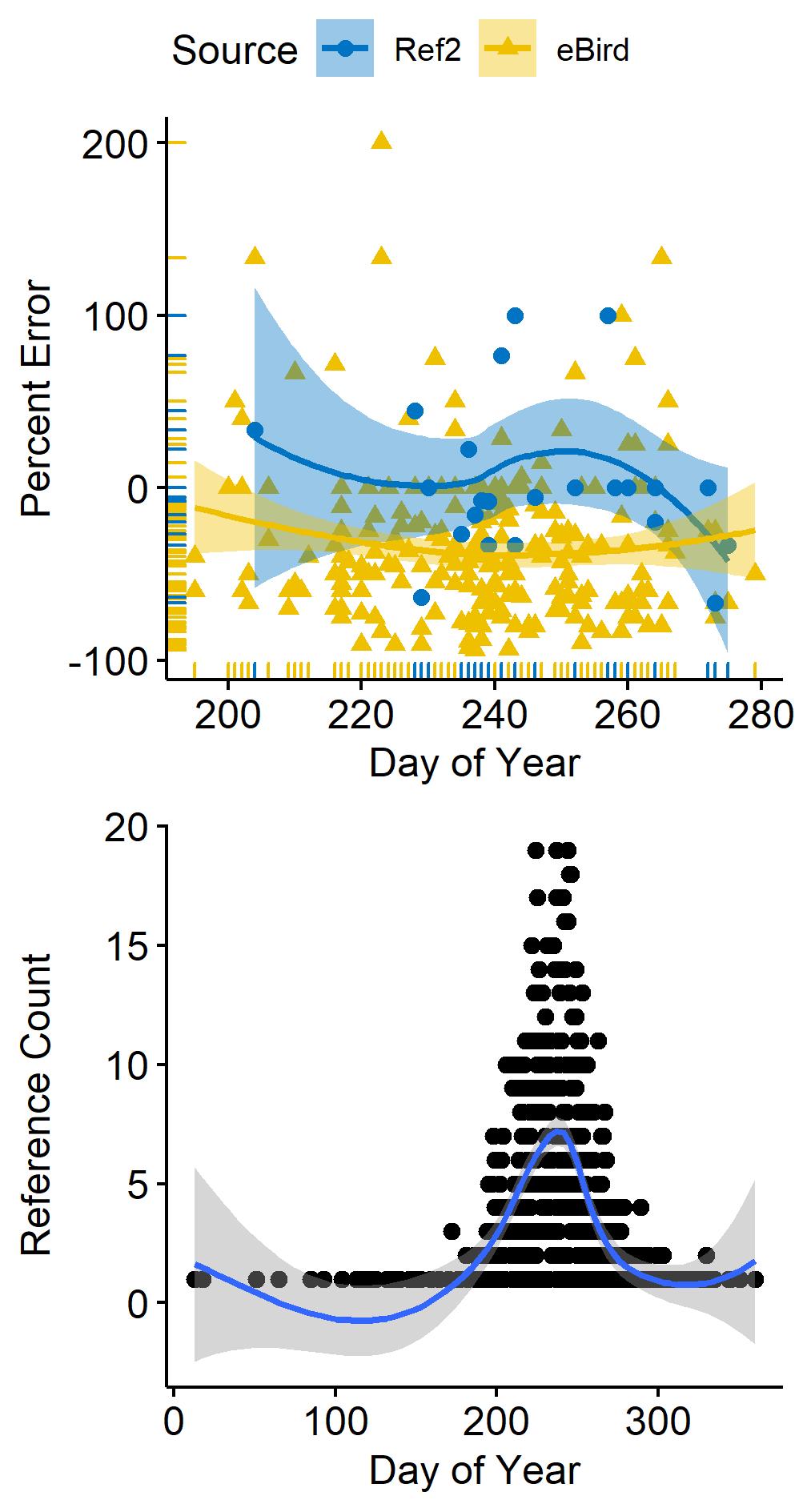

### RingbilledGull_Julian_PercentError.jpg

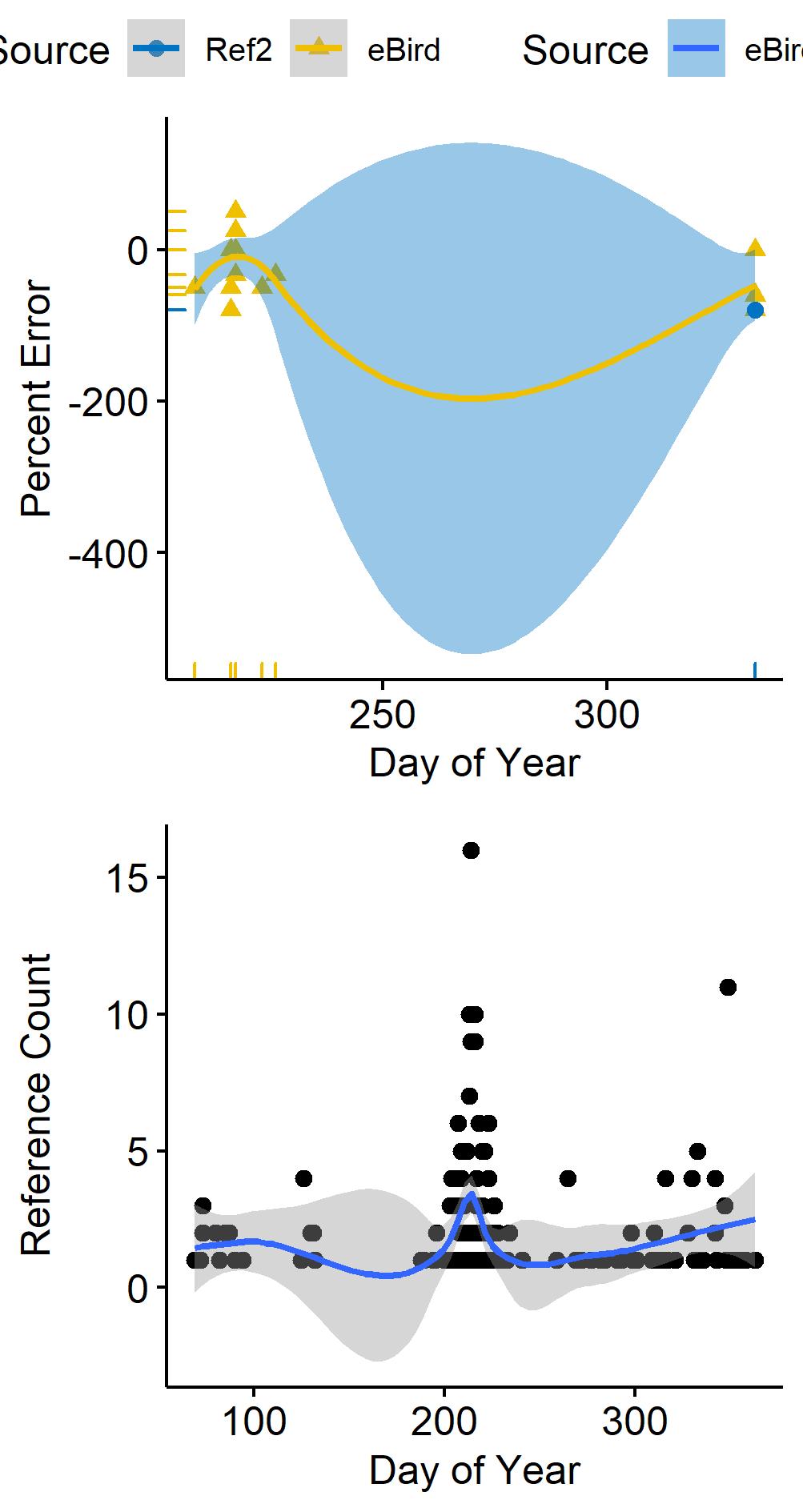

### RingneckedDuck_Julian_PercentError.jpg

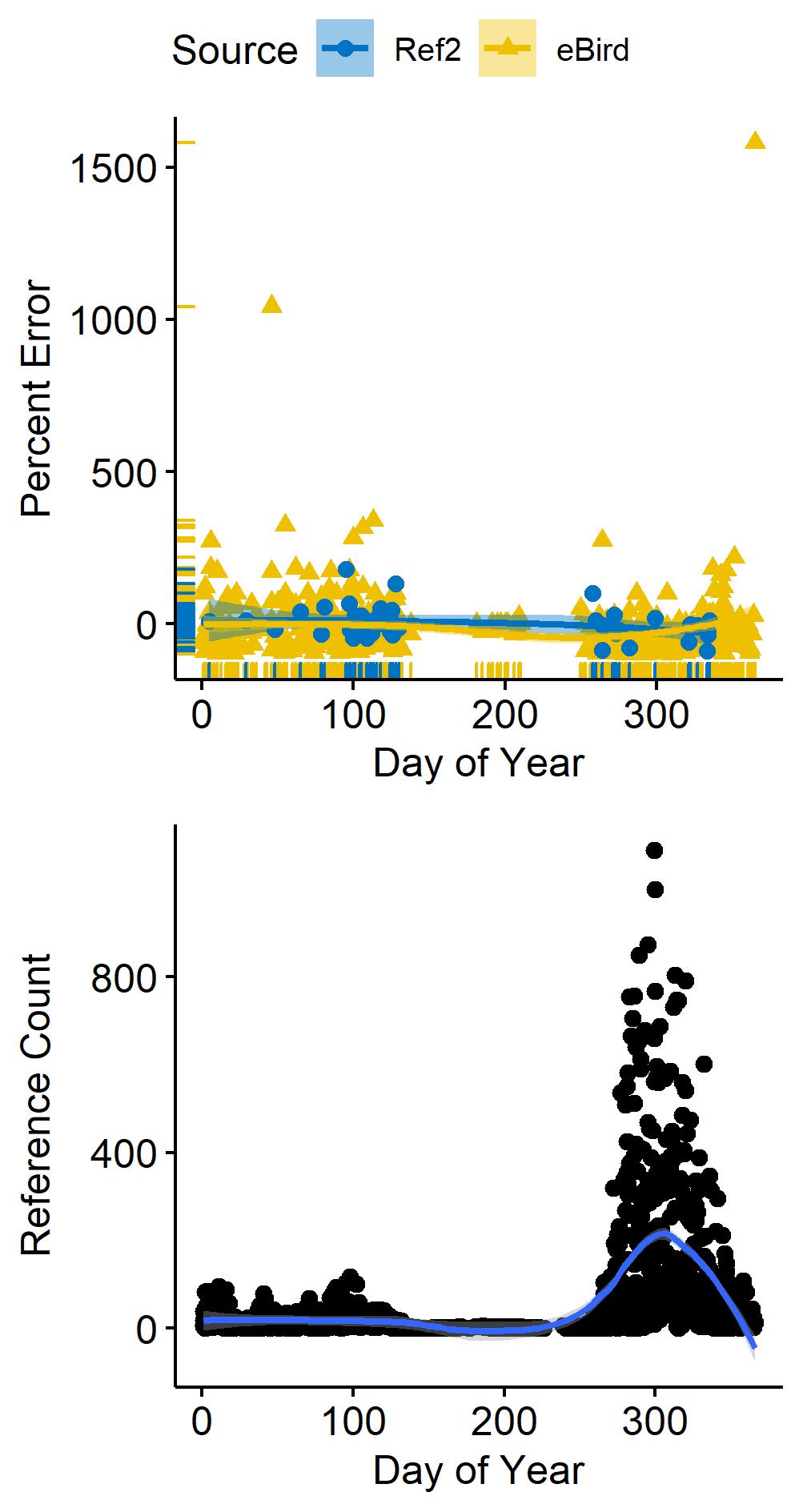

### RuddyDuck_Julian_PercentError.jpg

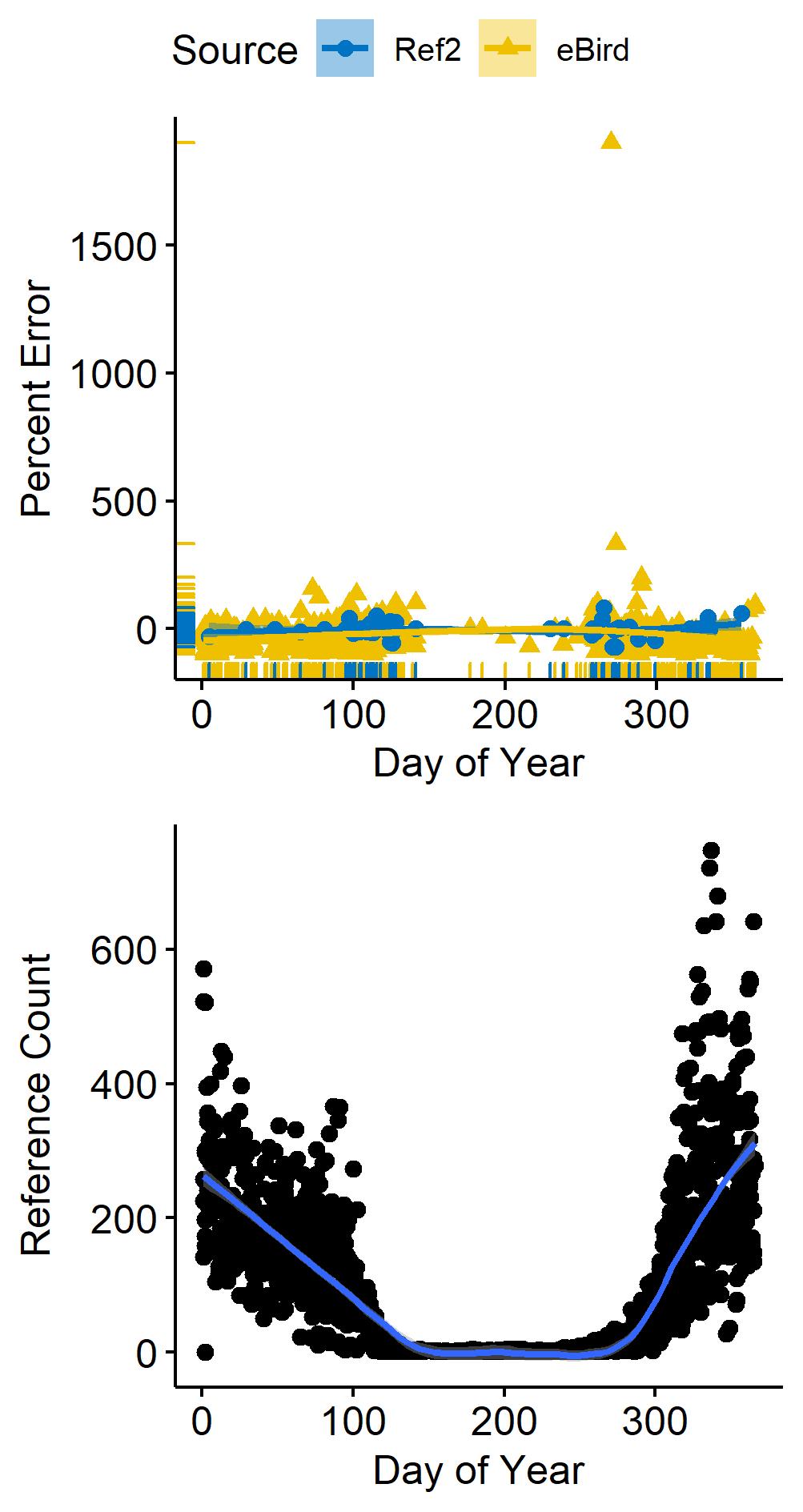

### SurfScoter_Julian_PercentError.jpg

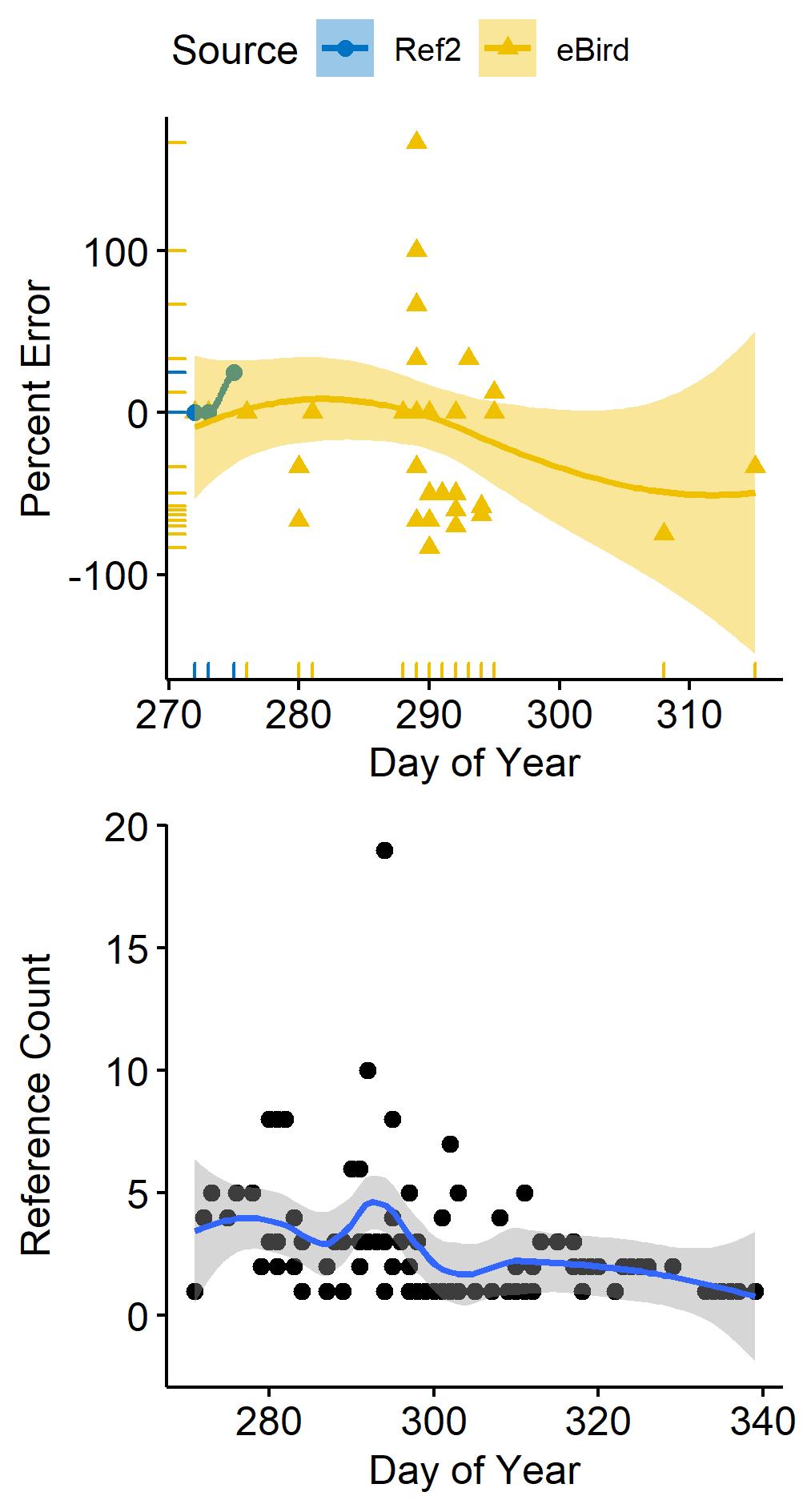

### WoodDuck_Julian_PercentError.jpg

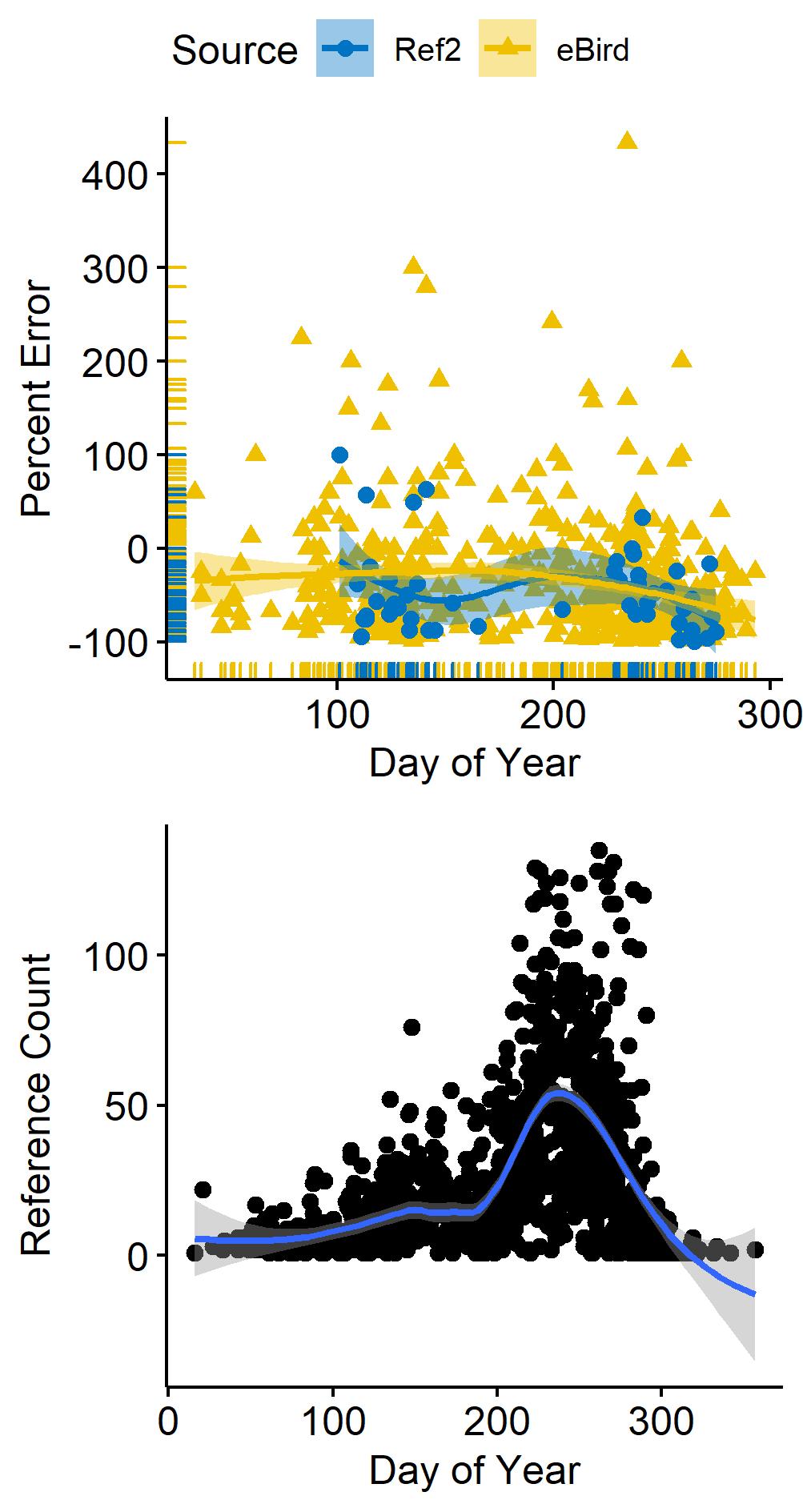
